# Microbial dysbiosis and metabolic disorders promote rheumatoid arthritis across successive stages: a multi-omics cohort study

**DOI:** 10.1101/2022.02.13.480248

**Authors:** Mingyue Cheng, Yan Zhao, Yazhou Cui, Chaofang Zhong, Yuguo Zha, Shufeng Li, Guangxiang Cao, Mian Li, Lei Zhang, Kang Ning, Jinxiang Han

## Abstract

**Background:** Rheumatoid arthritis (RA) is a chronic inflammatory disease that typically progresses through four stages, from mild stiffness to severe disability. The critical involvement of gut microbial dysbiosis in promoting RA has been intriguing. The aim of this study was to investigate stage-specific roles of microbial dysbiosis and metabolic disorders in pathogenesis across successive stages of RA.

**Methods:** In this multi-omics cohort study, fecal metagenomic, plasma metabolomic data were collected from 76 patients grouped into four RA stages, 19 patients with osteoarthritis, and 27 healthy individuals in China. A non-redundant gene catalogue was constructed, functionally annotated, and clustered into metagenomic species (MGS). Leave-one-out analysis was performed to determine the MGS that most drove the microbial dysfunctions with RA across stages. Random forest algorithm was performed to investigate diagnostic potential of microbial or metabolic features for each stage. Additionally, to verify the bacterial invasion of the joint synovial fluid, we conducted 16S rRNA gene sequencing, bacterial isolation, and scanning electron microscopy on joint synovial fluid from another cohort of 271 RA patients grouped into four RA stages.

**Findings:** We found that microbial dysbiosis and metabolic disorders had stage-specific effects on RA progression. For instance, arginine succinyltransferase pathway was elevated mainly in the second and third stages of RA (*p*=1·4×10^−3^), which was driven by *Escherichia coli*, and it was correlated with the increase of the rheumatoid factor (*p*=1·35×10^−3^). Considerable elevation of methoxyacetic acid (*p*=1·28×10^−8^) and cysteine-S-sulfate (*p*=4·66×10^−12^) might inhibit osteoblasts in the second stage and enhance osteoclasts in the third stage, respectively. Moreover, fecal microbes *Eggerthella lenta* and *Bifidobacterium longum* that were exclusively elevated in the fourth stage, were also detected in the joint synovial fluid.

**Interpretation:** Our findings elucidate for the first time the stage-specific roles of microbial dysbiosis and metabolic disorders across successive stages of RA, which open up new avenues for RA prognosis and therapy. We demonstrate the buildup of these effects might induce microbial invasion of the joint synovial fluid in the fourth stage of RA.

## Introduction

Rheumatoid arthritis (RA) affects over tens of millions of people worldwide.^1^ RA is recognized clinically as a chronic, inflammatory, autoimmune disease that primarily affects the joints and typically has four stages:^2–4^ 1) In the first stage, the synovium of the joints is inflamed and most people have minor symptoms such as stiffness upon awakening; 2) In the second stage, the inflamed synovium has caused damage to the joint cartilage and people begin to feel swelling, and have a restricted range of motion; 3) In the third stage, RA has proceeded to a severe state when bone erosion begins and the cartilage on the surface of the bones has deteriorated, resulting in the bones rubbing against one another; and 4) In the fourth stage, certain joints are severely deformed and lose function. To inhibit RA progression, specific therapeutic strategies are necessary for people across different RA stages.

Gut microbial dysbiosis has been implicated in the pathogenesis of RA via a range of mechanisms such as metabolic perturbation and immune response regulation, which is known as the gut-joint axis:^5^ For instance, increased abundance of *Prevotella* and *Collinsella* in RA patients are correlated with the production of TH17 cell cytokines.^6,7^ Metabolites have also been correlated to immunity regulation in RA: administration of Short-chain fatty acids to mice with collagen-induced arthritis (CIA) can reduce the severity of arthritis by modulation of IL-10.^5,8^ Comprehensive metagenomic and metabolomic analyses could therefore enhance our understanding about the gut-joint axis. However, the role of the gut-joint axis across successive stages of RA is understudied,^5,9^ where more examinations may provide an alternative approach to ameliorate RA progression.

Here, we aimed to investigate whether microbial dysbiosis and metabolic disorders had stage-specific effects in RA pathogenesis, and whether or in which stage microbial invasion of the joint synovial fluid happened.

## Methods

### Study design and participants

For the multi-omics study (**figure 1**), a total of 122 fecal and 122 plasma samples were collected from 122 outpatients of the Shandong Provincial Qianfoshan Hospital (Jinan, Shandong, China). These outpatients included 76 patients with RA, 19 patients with OA, and 27 healthy individuals (**Table**). RA patients were grouped into four RA stages including RAS1 (n=15), RAS2 (n=21), RAS3 (n=18), and RAS4 (n=22) according to the rheumatoid diagnostic score,^3^ where RAS1, RAS2, RAS3, and RAS4 has a score of 6–7, 8, 9, and 10, respectively. The score was evaluated by the sum of four categories as summarized in the 2010 rheumatoid arthritis classification criteria.^3^ Subsequently, fecal samples were sequenced and plasma samples were used to test the plasma metabolites, anti-citrullinated protein antibody, erythrocyte sedimentation rate, C-reactive protein, rheumatoid factor, and cytokines. Plasma metabolites were tested using UHPLC-MS/MS, while inflammatory cytokines were quantified using the MESO SCALE DISCOVERY Quick Plex S600MM multiplex assay.

**Figure 1:**
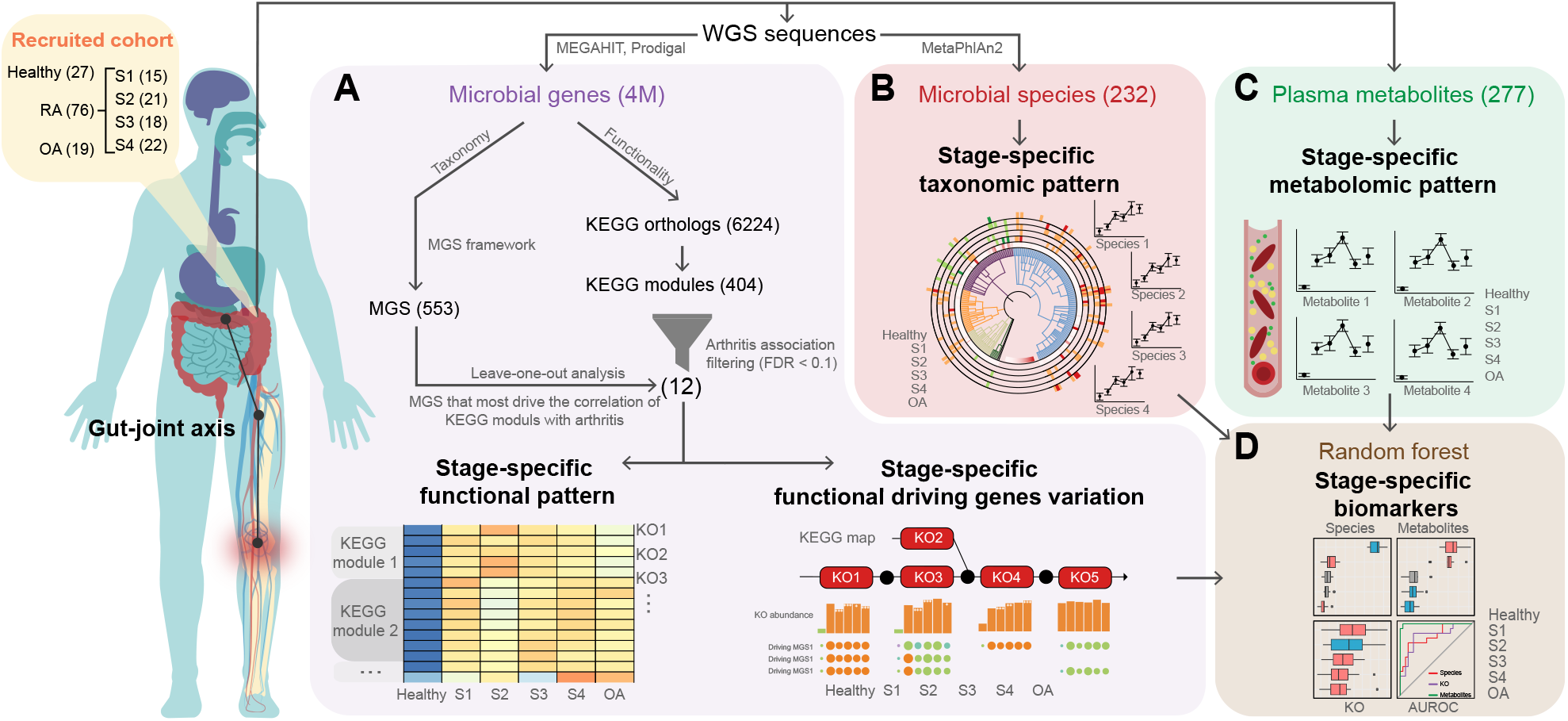
Schematic representation of the multi-omics study design. The multi-omics cohort included 76 outpatients with RA, 19 patients with OA, and 27 healthy individuals. (**A**) Microbial genes, predicted from WGS sequences, were clustered into MGS and mapped to KOs for functional annotation. The functional patterns across four successive stages, and the driven MGS of the functions were investigated. (**B**) A total of 232 classified microbial species were used to describe the taxonomic pattern across stages. (**C**) A total of 277 plasma metabolites were used to describe the metabolomic pattern across stages. (**d**) Random forest algorithm was performed on 6224 KOs, 232 microbial species, and 277 plasma metabolites from (A–C) to detect the stage-specific biomarkers at the multi-omics level. RA=rheumatoid arthritis. OA=osteoarthritis. MGS=metagenomic species. KO=KEGG ortholog.

All of the participants were at fasting status during the sample collection in the morning. Only participants who met the standard were recruited in this study: Recruited individuals had not received treatment in the recent month, and had no malignant tumor, no other rheumatic diseases such as ankylosing spondylitis, psoriasis, gout, no gastrointestinal diseases such as diarrhea, constipation, and hematochezia in the recent one month, no infections, no other comorbidity such as diabetes and hepatitis B.

To confirm the bacterial invasion of the joint synovial fluid, another cohort of 271 patients with RA of four distinct stages were recruited, including 52 RAS1, 66 RAS2, 67 RAS3, and 86 RAS4. RA patients were grouped into four RA stages according to the rheumatoid diagnostic score.^3^ The patients met the aforementioned standard. The entire experiment was conducted in a completely sterile atmosphere. For each patient, a total of 7ml synovial fluid was collected, of which 5ml was utilized for 16S rRNA gene sequencing, 1ml was used for bacteria isolation, and 1ml synovial fluid was prepared for scanning electron microscopy (**appendix p 1**).

The study was approved by the Ethics Committee of The First Affiliated Hospital of Shandong First Medical University (NO.2017-02).

### Procedures

#### Metagenome sequencing and processing

Whole-genome shotgun sequencing of fecal samples was carried out on the Illumina Hiseq X Ten. All samples were paired-end sequenced with a 150-bp read length. Reads containing more than 40 base pairs with low-quality values of 38 or less were discarded. Reads containing more than 10 unidentified base pairs were discarded. Reads containing more than 15 base pairs that overlapped adapter sequences were discarded. Next, reads were mapped against the human genome hg38 using Bowtie2^10^ (version 2·2·9). Those that were mapped were considered to be derived from the human genome and were discarded. Finally, unpaired reads were discarded. High-quality reads were used for the following analyses. MetaPhlAn2^11^ (version 2.7.8) was then used to generate microbial taxonomic profiles.

#### Contig assembly and ORF prediction

Reads were assembled into contigs using MEGAHIT^12^ (version 1·2·6) with the minimum contig length set at 500 bp and default parameters for metagenomics. The open reading frames (ORFs) were predicted from the assembled contigs using Prodigal^13^ (version 2·6·3) with default parameters. The ORFs of <100 bp were removed.

#### Non-redundant gene catalogue construction

The ORFs were clustered to remove redundancy using Cd-hit^14^ (version 4·6·6) with a sequence identity threshold set at 0·95 and the alignment coverage set at 0·9, which resulted in a catalogue of 4,047,645 nonredundant genes. The non-redundant genes were then collapsed into metagenomic species (MGS),^15,16^ and grouped into KEGG functional modules,^15^ as described below.

#### Identification of metagenomic species

High-quality reads were mapped to the catalogue of nonredundant genes using Bowtie 2^10^ (version 2·2·9) with default parameters. The abundance profile for each catalogue gene was calculated as the sum of uniquely mapped sequence reads, using 19M sequence reads per sample (downsized). The co-abundance clustering of the 4,047,645 genes was performed using canopy algorithm^16^ (http://git.dworzynski.eu/mgs-canopy-algorithm), and 553 gene clusters that met the previously described criteria^16^ and contained more than 700 genes were referred to as MGS. MGS that were present in at least 4 samples were used for the following analysis. The abundance profiles of MGS were determined as the medium gene abundance throughout the samples. MGS were taxonomically annotated by summing up the taxonomical annotation of their genes as described by Nielsen *et al*.^16^ Each MGS gene was annotated by sequence similarity to the bacterial complete genome in NCBI Reference Sequence Database (BLASTN, E-value<0·001).

#### Annotation of KEGG modules

The catalogue of the nonredundant genes was functionally annotated to the KEGG database (release 94.0) by KofamKOALA (version 1·3·0).^17^ The produced KEGG Orthologs (KOs) were mapped to the KEGG modules annotation downloaded on 1 August 2020 from the KEGG BRITE database. KOs that were present in at least 4 samples were used for the following analysis. The KO abundance profile was calculated by summing the abundance of genes that were annotated to each of the KOs.

#### Leave-one-out analysis

Leave-one-out analysis was used to test which MGS was driving the observed associations between KEGG modules and arthritis. The calculation of the KO abundance was iterated excluding the genes from a different MGS in each iteration. The effect of a given MGS on a specified association was defined as the change in median Spearman correlation coefficient between KOs and arthritis when genes from the respective MGS were left out, as previously described.^15^

#### Multi-omics biomarkers

Random forest algorithm was performed using the R package “randomForest”. Function “trainControl” in R package “caret” was used to perform 10 repeats of 10-fold cross-validation for each data set. Function “train” in R package “caret” was used to fit models over different tuning parameters to determine the “mtry” for random forest algorithm.

#### Correlation of phenotypes and KEGG modules

The clinical phenotypes including types of arthritis (Healthy=0, OA=1, RA=2) and plasma levels of cytokines were used in the correlation analysis with KEGG modules. The associations of clinical phenotypes with KEGG modules were determined by if Spearman correlations of the phenotype with the abundances of KOs in the given KEGG module were significantly higher or lower (Mann–Whitney U-test FDR<0·1) than with the abundances of all other KOs. The phenotypes adjusted by age and gender were also tested using partial Spearman correlation. The union set of the significant associations between KEGG modules and phenotypes/phenotypes adjusted by age and gender were used for the following analysis.

### Statistical analysis

The correlations between vectors such as microbial species, metabolites, and the levels of cytokines, were tested using Spearman correlation or partial Spearman correlation with age and gender adjusted (*P* values), corrected by FDR (q values). For categorical metadata, samples were pooled into bins (RA/OA/Healthy, RAS1/RAS2/RAS3/RAS4) and significant features were identified using Mann-Whitney-Wilcoxon Test (*P* values) with Benjamini and Hochberg correction (FDR, q values).

### Role of the funding source

The funders of the study had no role in study design, data collection, data analysis, data interpretation, or writing of the report.

## Results

We obtained a total of 231 classified microbial species from metagenomic data, and tested their alterations in each stage of RA, as compared to healthy controls (**figure 2**). The elevated species in RA progression were mostly from the phyla Firmicutes and Actinobacteria, while the depleted species were predominantly from the phylum Bacteroides. In addition to 15 microbial species that were found substantially altered in RA or OA (*q*<0·05, **appendix p 3**), other previously reported RA-related species were specified in boxplots (**figure 2**).^18^ We found although a few microbes were identified as correlated with RA, they might not remain plentiful or depleted throughout the progression. *Bifidobacterium dentium*, for instance, was reported to be associated with the development of dental caries and periodontal disease, both of which were particularly prevalent in patients with RA.^19,20^ Compared to healthy controls, it maintained considerably elevated across RA stages except for RAS1 (RAS2: *p*=7·16×10^−3^, RAS3: *p*=3·70×10^−3^, RAS4: *p=*9·15×10^−4^).

**Figure 2:**
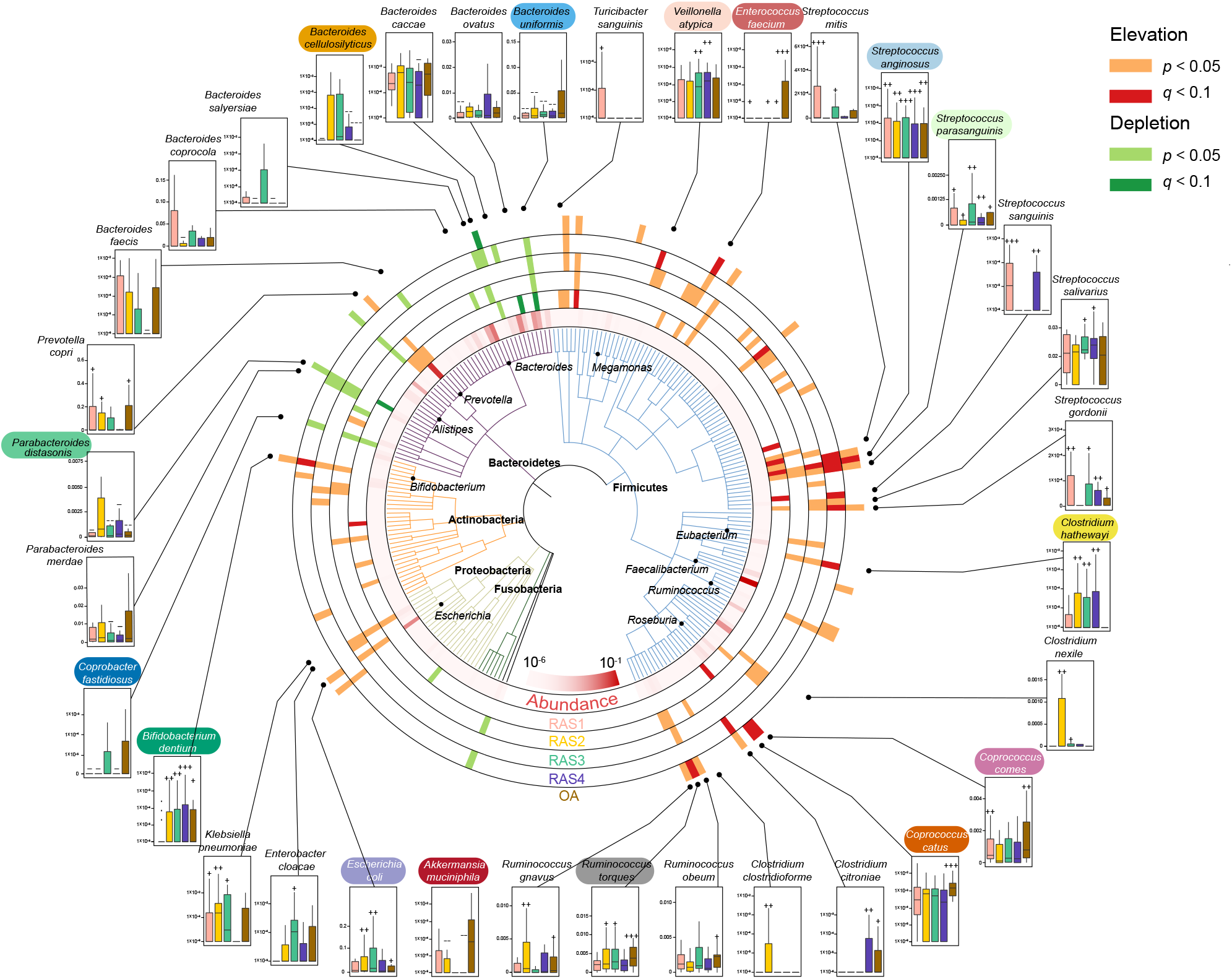
Microbial taxonomic patterns across four stages of RA. In total, 232 classified species whose relative abundance was more than 1×10^−6^ are shown in a phylogenetic tree, grouped in the phyla Fusobacteria, Proteobacteria, Actinobacteria, Bacteroidetes, and Firmicutes. In the outer circles, species at each arthritis stage are marked for significant elevation (*p*<0·05, orange; *q*<0·1, red) or depletion (*p*<0·05, light green; *q*<0·1, dark green) in abundance, as compared to those of healthy individuals. The innermost circle shows species relative abundances averaged over all samples. Box plots show the relative abundances of 15 species identified as notably altered in RA or OA (name in color; *q*<0·05; **appendix p 2**) and others that have been correlated with RA in previous studies.^18^ The order and color of the box are consistent with the stage circle. Significant changes are denoted as follows: +++, elevation with *p*<0·005; ++, elevation with *p*<0·01; +, elevation with *p*<0·05; –––, depletion with *p*<0·005; ––, depletion with *p*<0·01; –, depletion at *p*<0·05; Mann-Whitney-Wilcoxon test. Boxes represent the interquartile range between the first and third quartiles and the line inside represents the median. Whiskers denote the lowest and highest values within the 1·5×interquartile range from the first and third quartiles, respectively. –

Inspired by this finding, we then investigated 29 species that were altered exclusively in a specific stage (*p*<0·05; **appendix pp 4–5**). We found that *Collinsella aerofaciens* was elevated exclusively in RAS1 (*p=*0·043). *C. aerofaciens* was previously reported to generate severe arthritis when inoculated into CIA-susceptible mice, and an *in vitro* experiment showed that *C. aerofaciens* could increase gut permeability and induce IL-17A expression, a key cytokine involved in RA pathogenesis.^7^ The elevation of *C. aerofaciens* in RAS1 might contribute to the early breach in gut barrier integrity, through which the translocation of microbial products would then trigger the subsequent clinical arthritis.^5^ Moreover, *Veillonella parvula*, whose infection could cause osteomyelitis,^21^ was found elevated exclusively in RAS3 (*p*=0·027). *Eggerthella lenta* (*p*=0·018) and *Bifidobacterium longum* (*p*=0·022) were found elevated exclusively in RAS4. The gavage of *E. lenta* were reported to increase gut permeability and produce proinflammatory cytokines IL-6, IL-21, IL-23.^22^ We also recognized species altered exclusively in OA, such as elevated *Dialister invisus* (*p*=0·041) that was positively correlated with spondyloarthritis severity.^23^ These stage-specific altered species had the potential to serve as the targets for intervention in a given RA stage.

Next, we sought to detect the microbial dysfunction across stages of RA. We grouped 4,047,645 metagenomic genes into 6,224 KOs and 404 KEGG modules. We identified 12 KEGG modules that were significantly correlated with RA or OA (*q*<0·1; **appendix p 6**), and presented their variation across stages (**figure 3A**). We then used leave-one-out analysis to identify the MGS that most drove the correlations of these KEGG modules with RA or OA. (**figure 3B; appendix p 7**).

**Figure 3.**
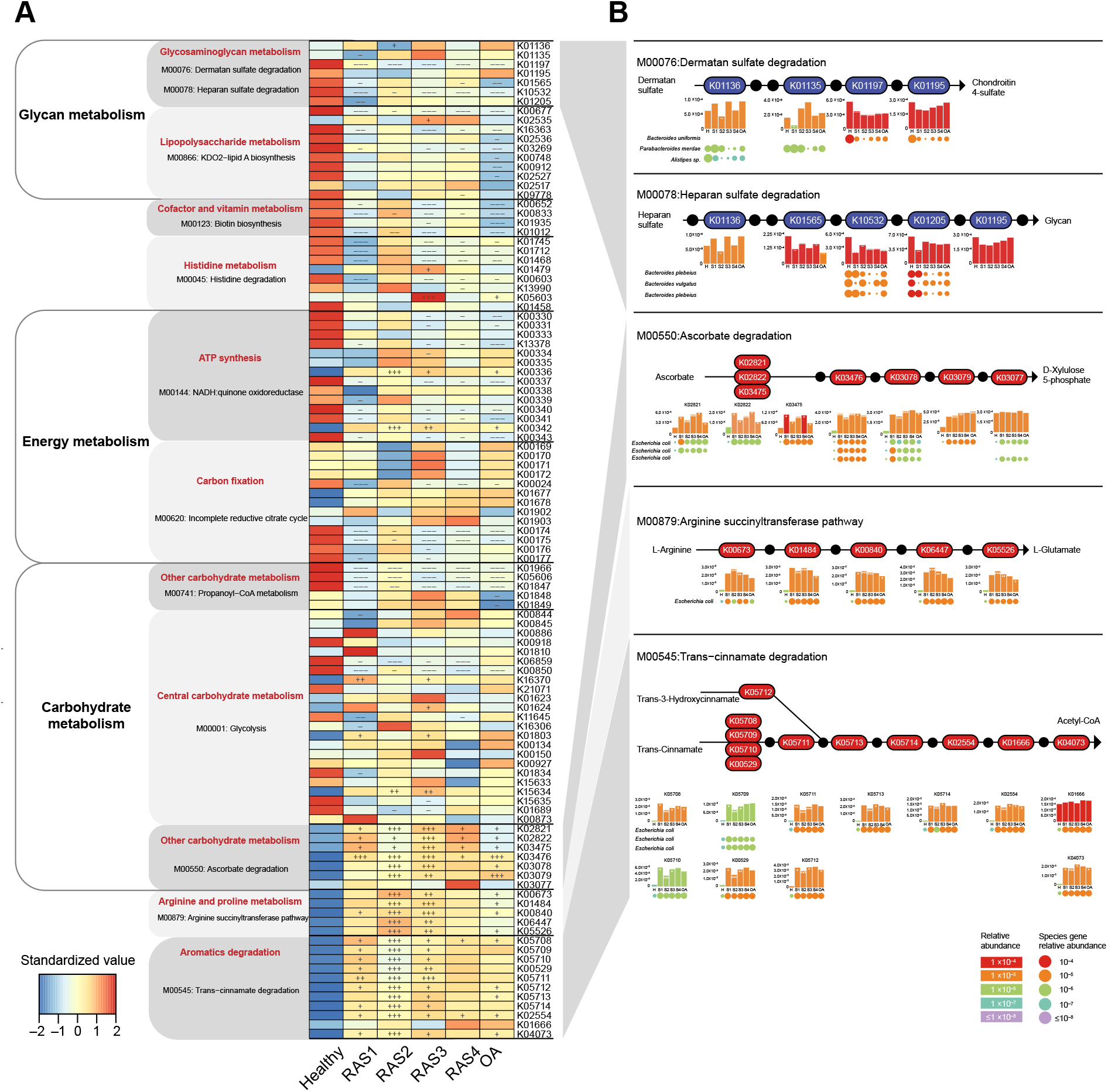
Microbial dysfunction across four stages of RA. Gene abundances were assessed for elevation or depletion in each of the arthritis stages, RAS1 (n=15), RAS2 (n=21), RAS3 (n=18), RAS4 (n=22), and OA (n=19) compared to the healthy individuals (n=27). (**A**) Relative abundance of KO genes in the KEGG modules that were significantly correlated with arthritis (*q*<0·1; **appendix p 5**). KO genes with a prevalence of 5% or higher are shown. (**B**) KO genes involved in specific KEGG pathway modules in (A) are shown in the KEGG pathway maps. Each box in a pathway represents a KO gene and is marked in red for elevation or in blue for depletion at any of the stages compared to healthy individuals. Bar plots show relative gene abundances averaged over samples within each of the five groups (healthy (H), RAS1 (S1), RAS2 (S2), RAS3 (S3), RAS4 (S4), and OA) and are colored according to the values. Each KO gene is composed of MGS genes represented by circles. The sizes and colors of the circles are proportional to the relative abundances of the MGS genes. MGS genes are grouped into one row and indicated by the taxonomic name. The three MGS that most drove the correlation of the KEGG modules with arthritis types are shown. In all panels, significant changes are denoted as follows: +++, elevation with *p*<0·005; ++, elevation with *p*<0·01; +, elevation with *p*<0·05; –––, depletion with *p*<0·005; ––, depletion with *p*<0·01; –, depletion at *p*<0·05; Mann-Whitney-Wilcoxon test. KO=KEGG ortholog. MGS=metagenomic species.

We found an evident decrease in glycosaminoglycan (CAG) metabolism across four RA stages and OA. It was mainly reflected by the significant decrease in K01197 (hyaluronoglucosaminidase) of dermatan sulfate (DS) degradation and the significant decrease in K10532 (heparan-alpha-glucosaminide N-acetyltransferase) of heparan sulfate (HS) degradation (*p*<0.05; **figure 3A**). Chondroitin 4-sulfate is a major component of the extracellular matrix of many connective tissues, such as cartilage, bone, and skin.^24^ We found that the significant depletion of DS degradation would inhibit the production of chondroitin 4-sulfate (**figure 3B**), which might hurt the mechanical properties of the articular cartilage.^24^ Moreover, the significant depletion of HS degradation might be a potential cause of the higher plasma level of HS observed in RA and OA patients,^25,26^ which could promote arthritis progression by regulating protease activity.^27^ The most driving species of DS degradation and HS degradation were MGS *Bacteroides uniformis* and MGS *Bacteroides plebeius*, respectively. The genes of MGS *B. uniformis* related to K01197 were found most depleted in RAS2, while the genes of MGS *B. plebeius* related to K10532 were found most depleted in RAS3 and RAS4 (**figure 3B**). These results indicated that the depleted microbial function in DS degradation and HS degradation driven by *B. uniformis* and *B. plebeius*, respectively, could promote RA and OA in a way of hurting articular cartilage.

We also identified elevated microbial functions that were related to inflammation. Ascorbate could prevent the development of inflammatory arthritis by facilitating collagen synthesis, moderating autoimmune responses, and ameliorating inflammation.^28^ Here, we found most of the KOs related to ascorbate degradation retained a higher level across RA stages and OA, especially in RAS2 and RAS3 (*p*<0.05; **figure 3A**). Genes of K02821 (phosphotransferase system) in RAS1, K03475 (phosphotransferase system), K03476 (L-ascorbate 6-phosphate lactonase), and K03479 (L-ribulose-5-phosphate 3-epimerase) were mostly contributed by MGS *Escherichia coli* (**figure 3B**). The enhanced ascorbate degradation might contribute to the deficiency of the ascorbate reported in patients with RA^29^ and were found positively correlated with multiple plasma cytokines (*q*<0·1; **appendix p 8**), such as IL-1β (*p*=5·44×10^−4^), TNF-α (*p*=6·59×10^−4^), and IL-6 (*p*=1·12×10^−3^). Moreover, to confirm the effects of ascorbate on RA progression, we examined the plasma TNF-α level and IL-6 level, bone CT scans, and bone density of 1) normal DBA/1 mice, 2) DBA/1 mice with CIA, and 3) DBA/1 mice with CIA and gavage of ascorbate. We found that the three-month gavage of ascorbate to CIA mice can prevent the increase of TNF-α and IL-6 levels by half, inhibit bone destruction, and maintain bone density (1·58 ± 0·0034 g/cm^3^), as compared to the CIA mice without ascorbate (1·53 ± 0·013 g/cm^3^), and the normal group (1·61 ± 0·021 g/cm^3^; **appendix p 2**,**9**).

For other elevated microbial functions, the trans-cinnamate degradation driven by MGS *E. coli*, where most KOs were notably elevated in RAS2, was also correlated with multiple cytokines (q<0·1; **appendix p 8**), such as IL-13 (*p*=1.63×10^−5^), IL-1β (*p*=2.87×10^−5^), and IL10 (*p*=4.10×10^−5^). Moreover, the arginine succinyltransferase pathway driven by MGS *E. coli* was found significantly elevated mainly in RAS2 and RAS3 (**figure 3**). L-arginine is able to prevent bone loss induced by zinc oxide nanoparticles or by cyclosporin A, through anti-inflammatory mechanism^30^ or nitric oxide production, respectively.^31^ Both arginine succinyltransferase pathway and trans-cinnamate degradation was positively correlated with the elevation of rheumatoid factor (*p*=1·35×10^−3^). Taken together, these results suggested that microbial dysfunction could promote RA progression mainly by hurting bone tissue and strengthening inflammation. The inflammation-related microbial dysfunction was extremely active in RAS2 and RAS3 and largely driven by *E. coli*.

Next, we investigated whether or in which stage microbial invasion of the joint synovial fluid happened. Enhanced gut permeability may render it possible for microbes and their products to translocate, triggering an immune response.^5,32^ We thus speculated that gut microbes might invade the joint synovial fluid of patients with RA through the gut-joint axis. To test this, we performed 16S rRNA gene sequencing on the synovial fluid samples from another cohort of 271 patients in four RA stages (**appendix p 1**), including RAS1 (n=52), RAS2 (n= 66), RAS3 (n=67), and RAS4 (n=86). Notably, we were not able to obtain enough bacterial DNA for sequencing in samples of RAS1, RAS2, or RAS3, however, we could identify many microbes in samples of RAS4 (**appendix p 10**). We found that most of the microbes in joint synovial fluid were from phyla Proteobacteria and Firmicutes. Moreover, we could recognize *Eggerthella lenta* and *Bifidobacterium longum* in most of the synovial fluid samples (**appendix p 10**), both of which were observed to be exclusively elevated in fecal metagenome of patients in RAS4 from the multi-omics cohort (**appendix p 5**). In addition, *Prevotella copri* that has been reported highly correlated with RA^6,33^ was also found abundant in most synovial fluid samples of patients in RAS4. We then randomly selected six synovial fluid samples per RA stage for bacteria isolation. Only from three synovial fluid samples of RAS4 can we separate bacteria. We then picked and sequenced three single colonies per synovial fluid sample. Five of the nine colonies were identified as *Clostridium sp*., three were identified as *Enterococcus sp*., and one was identified as *Citrobacter sp*. (**appendix p 11**). We subsequently observed the corresponding synovial fluid samples using scanning electron microscopy, and found substances shaped like bacteria in rod-like or spherical forms (**appendix p 12**). Taken together, this multi-faceted investigation has provided unprecedented evidence to support the existence of microbial invasion of the joints in the fourth stage of RA.

We then introduced metabolomic data, and performed a random forest algorithm on 232 microbiome species, 6224 KOs, and 277 metabolites to test their diagnostic potential for each stage of RA and OA (**figure 4A–E**). Metabolites exhibited the best AUROC in discriminating samples of four RA stages or OA from healthy samples, with AUROC ranging from 0·974 to 0·998. Other characteristics at the species and KO levels exhibited weaker discriminant ability than features at the metabolite level, with AUROC ranging from 0·760 to 0·838 and from 0·799 to 0·852, respectively. The predominant metabolic disorders implicated a critical involvement in pathogenesis and a great diagnostic potential for RA stages. The most prominent changes in metabolites were the significant increase of DL-lactate and gly-glu in RAS1 (*p*=2·15×10^−6^, *p=*4·70×10^−4^), the decrease of N,N-dimethylaniline and the increase of methoxyacetic acid in RAS2 (*p*=4·60×10^−8^, *p*=1·28×10^−8^), the increase of cysteine-S-sulfate (*p*=4·66×10^−12^) in RAS3, the increase of galactinol and 3α-mannobiose in RAS4 (*p=*5·71×10^−5^, *p=*5·68×10^−4^), and the decrease of N,N-dimethylaniline and increase of gly-glu (*p=*2·00×10^−7^, *p=*1·74×10^−3^) in OA, as compared to a healthy state.

**Figure 4.**
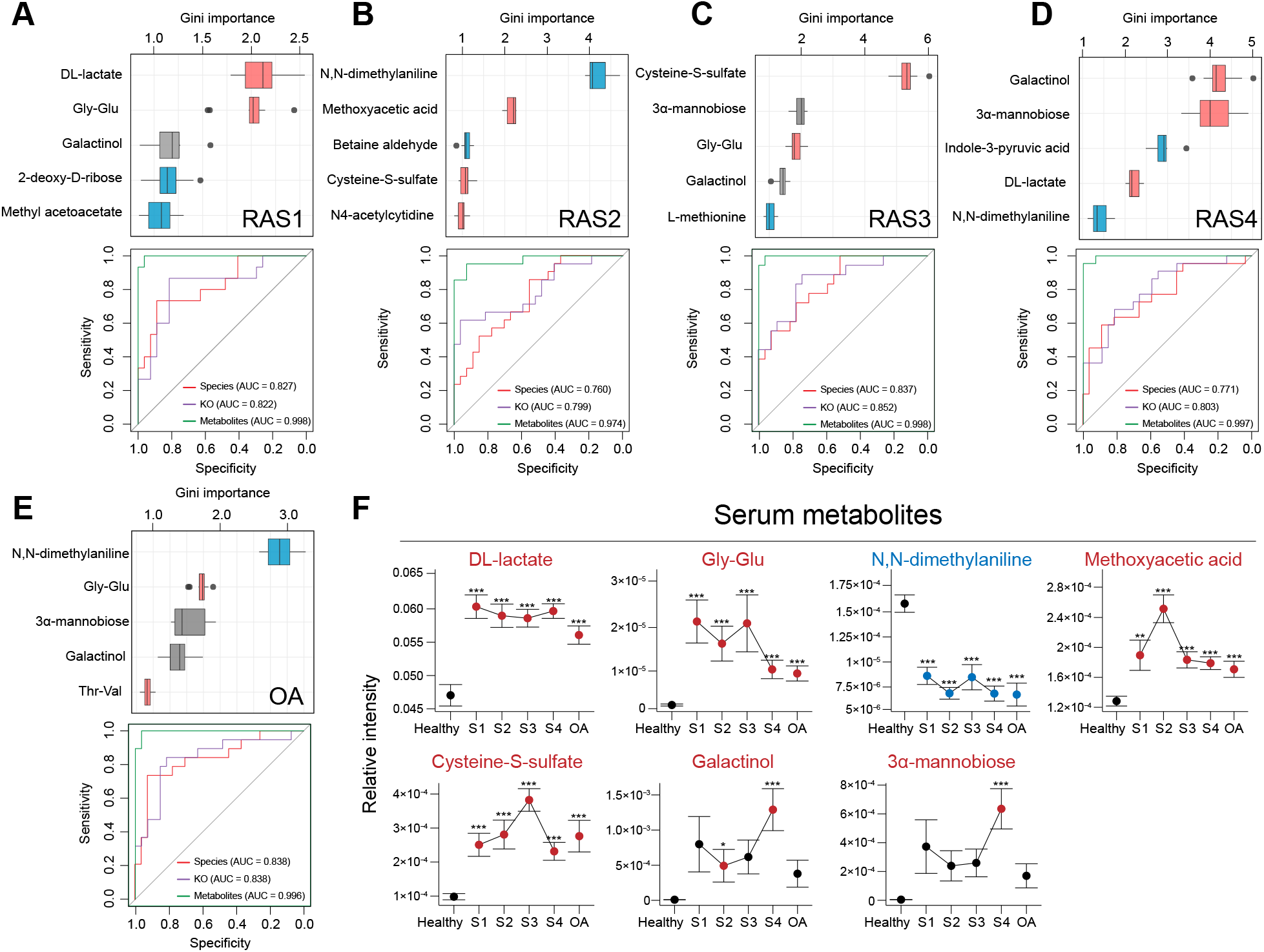
Multi-omics diagnostic potential for the RA stage. A random forest algorithm was performed on 6224 KOs, 232 microbial species, and 277 plasma metabolites in RAS1 (**A**), RAS2 (**B**), RAS3 (**C**), RAS4 (**D**), and OA (**E**). The Gini importance of the top five most discriminant metabolites are displayed. Boxes represent the interquartile range between the first and third quartiles and the line inside represents the median. Whiskers denote the lowest and highest values within the 1·5×interquartile range from the first and third quartiles, respectively. Boxes are marked in a specific color to show the significant elevation (*p*<0·05, red, Mann-Whitney-Wilcoxon test) or depletion (*p*<0·05, blue, Mann-Whitney-Wilcoxon test) of the features in each of the arthritis stages compared to the healthy group. The ROC curves of the random-forest model using microbial species, KOs, or metabolites were plotted, with AUC calculated by 10 randomized 10-fold cross-validation. The color of the curve represents the category of the used features. (**F**) The dot plots show stage-specific abundance or concentration (mean ± s.e.m.) of plasma metabolites, which are specified in (A–E). Four RA stages are connected to display the variance. Dots are colored differently if the features are significantly elevated (red) or significantly depleted (blue), as compared to those of the healthy group. ***, *P*<0·005; **, *P*<0·01; *, *P*<0·05; Mann-Whitney-Wilcoxon test. KO=KEGG ortholog

Moreover, metabolic disorders could distinguish a given RA stage from not just healthy controls but also other RA stages or OA (**figure 4F**): Methoxyacetic acid in RAS2 (*p*=1·68×10^−4^) or cysteine-S-sulfate in RAS3 (*p*=2·42×10^−4^) or Galactinol and 3α-mannobiose in RAS4 (*p*=9·37×10^−3^, *p*=4·89×10^−3^), respectively, was higher than that in all the other RA stages and OA. In addition, DL-lactate in OA was less than that in all RA stages (*p*=0·037), which might improve clinical differentiation of early RA from OA.

## Discussion

Our findings reveal dynamic shifts in gut microbiome and plasma metabolome, and their continuous roles in pathogenesis of RA across four successive stages (**figure 5**). Moreover, we demonstrate the buildup of these effects might induce microbial invasion of the joint synovial fluid in the last stage of RA.

**Figure 5.**
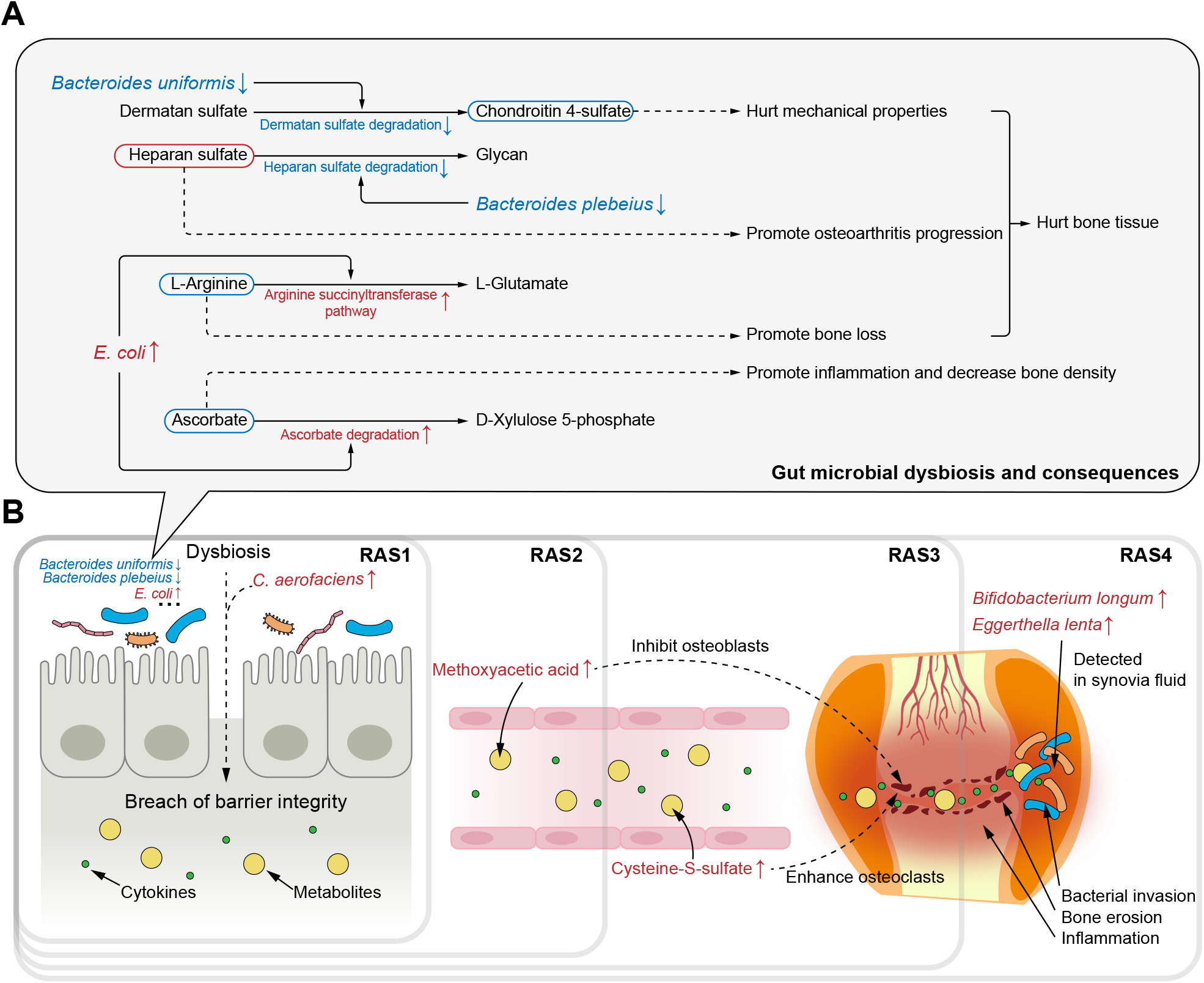
Potential pathogenesis across successive RA stages from multi-omics perspective. (**A**) Potential mechanisms by which gut microbial dysbiosis play roles in RA pathogenesis through hurting hone tissue and increasing inflammation. The driving species, microbial dysfunction, and related metabolites were extracted from **figure 1B**. The red or blue box of metabolites represents their speculated elevation or depletion according to the KEGG map. The dotted line represents the speculated effects of microbial and metabolic variation on arthritis pathogenesis (**B**) The most representative effects of microbial dysbiosis and metabolic disorders on RA progression across successive stages.

Clinical microbial intervention should take into account the stage of RA. Previous studies mainly focused on the preclinical stage or did not consider the stage.^5,9^ However, we found each RA stage had its special elevated or depleted microbes that played a role in RA pathogenesis (**figure 5A**). Hence, it may not be adequate for clinical guidance to generally report microbial alterations in RA without information of the stage. For instance, early inhibition of *Collinsella aerofaciens* that was elevated exclusively in the first stage could help prevent increasing of gut permeability.^7^ Additionally, inhibition of *Escherichia coli* in the second and third stage could help maintain the content of L-arginine that acted as an inhibitor of bone lose,^30,31^ as well as the content of anti-inflammatory ascorbate.^28^ Moreover, certain species may need intervention across stages owing to its depletion during the whole RA progression. A cross-stages restoration of *Bacteroides uniformis* could help maintain the content of chondroitin 4-sulfate to keep mechanical properties of the articular cartilage.^24^

Moreover, metabolic alterations kept considerable throughout RA progression, in spite of which we found that certain of these metabolites need a higher priority of intervention in a specific stage. In the second stage of RA, the aberrant elevation of methoxyacetic acid might have inhibitory effects on osteoblasts and cause reductions in bone marrow cellularity^34–36^ (**figure 5B**). The inhibited osteoblasts then drew the foreshadowing for the bone erosion that happened in the next stage. In the third stage, the considerable elevation of cysteine-S-sulfate might enhance the osteoclasts by NMDA-R interaction^37^. The imbalance between osteoblasts and osteoclasts would then promote bone erosion that happened in the third stage and persisted in the late RA stages. Thus, methoxyacetic acid may be a targeted metabolite for treatment to patients in the second stage of RA and serve as a precaution against the upcoming third stage.

Our findings suggested that bacterial invasion of joint synovial fluid happened in the fourth stage of RA (**figure 5B**). Joint synovial fluid was generally considered sterile, and indeed, we failed to either extract enough DNA or isolate bacteria from the synovial fluid in the first three stages. However, in the fourth stage, we succeeded to obtain bacterial 16S reads, isolate bacteria, and observe substances shaped like bacteria in rod-like or spherical forms under scanning electron microscopy. Moreover, in the multi-omics cohort, we found two fecal microbes elevated exclusively in the fourth stage of RA, *Eggerthella lenta* and *Bifidobacterium longum*, and their existence in the joint synovial fluid was validated by the other cohort. It might due to the buildup of the continuous damages in gut barrier and microbes and microbial metabolites would then be transferred to the joints via blood.^5^ Hence, for patients in the fourth stage of RA, in addition to routine medical therapies, specific treatments to the microbes in the joint synovial fluid may ameliorate the joint micro-environment to decrease synovial inflammation and inhibit potential bacterial effects.

This study also has limitations and prospects. Firstly, a long-term follow-up investigation on a single individual throughout his/her RA development may reinforce the conclusions of this study. Secondly, it remains unclear how bacterial genetic materials are transferred from intestine to joint. It might be realized by bacteria transmission through blood or by means of extracellular vesicles or both. Thirdly, the proposed links between microbial dysbiosis/metabolic disorders and RA can serve as a guidance for future experiments on RA pathogenesis. Lastly, additional researches into the synovial fluid microbiome and metabolome have the potential to reveal more sophisticated mechanisms underlying RA pathogenesis.

In conclusion, this study demonstrates microbial and metabolic roles in RA pathogenesis across four successive stages. A stage-specific intervention of microbial dysbiosis and metabolic disorders is warranted for prognosis and prevention of RA.

## Contributors

MC, YZ, LZ, KN, and JH planned the study, reviewed and verified the data. MC, YZ, YC, CZ, YZha, SL, GC, and ML collected samples and conducted experiments. MC, YZ, and KN conducted data analysis and produced the figures and tables. MC, YZ, KN, and JH wrote the manuscript. All authors revised the manuscript. MC, YZ, LZ, KN, and JH supervised the study. All authors had full access to all the data in the study and had final responsibility for the decision to submit for publication.

## Declaration of interests

We declare no competing interests.

## Data sharing

Whole-genome shotgun sequencing data are available in the Genome Sequence Archive (GSA) section of the National Genomics Data Center (project accession number CRA004348). 16S rRNA gene sequencing data are available in the Genome Sequence Archive (GSA) section of the National Genomics Data Center (project accession number CRA005811).

## Acknowledgments

This work was partially funded by the National Natural Science Foundation of China (grant numbers 31871334, 82003766, 32071465, and 31671374), the Academic Promotion Programme of Shandong First Medical University (grant number 2019LJ001), and the Key Research and Development project of Shandong Province (grant number 2021ZDSYS27).

## Notes

### Competing Interest Statement

The authors have declared no competing interest.

### Summary of Updates

We have revised some typos in the manuscript. We have reorganized the construction of the manuscript.

## References

1. Almutairi K, Nossent J, Preen D, Keen H, Inderjeeth C. The global prevalence of rheumatoid arthritis: a meta-analysis based on a systematic review. Rheumatol Int 2021; 41(5): 863–77.

2. Smolen JS, Aletaha D, McInnes IB. Rheumatoid arthritis. Lancet 2016; 388(10055): 2023–38.

3. Aletaha D, Neogi T, Silman AJ, et al. 2010 Rheumatoid arthritis classification criteria: an American College of Rheumatology/European League Against Rheumatism collaborative initiative. Arthritis Rheumatol 2010; 62(9): 2569–81.

4. Steinbrocker O, Traeger CH, Batterman RC. Therapeutic criteria in rheumatoid arthritis. J Am Med Assoc 1949; 140(8): 659–62.

5. Zaiss MM, Joyce Wu HJ, Mauro D, Schett G, Ciccia F. The gut-joint axis in rheumatoid arthritis. Nat Rev Rheumatol 2021; 17(4): 224–37.

6. Pianta A, Arvikar S, Strle K, et al. Evidence of the immune relevance of Prevotella copri, a gut Microbe, in patients with rheumatoid arthritis. Arthritis Rheumatol 2017; 69(5): 964–75.

7. Chen J, Wright K, Davis JM, et al. An expansion of rare lineage intestinal microbes characterizes rheumatoid arthritis. Genome Med 2016; 8(1): 43.

8. Smith PM, Howitt MR, Panikov N, et al. The microbial metabolites, short-chain fatty acids, regulate colonic Treg cell homeostasis. Science 2013; 341(6145): 569–73.

9. Maeda Y, Takeda K. Role of gut microbiota in rheumatoid arthritis. J Clin Med 2017; 6(6): 60.

10. Langmead B, Salzberg SL. Fast gapped-read alignment with Bowtie 2. Nat Methods 2012; 9(4): 357–9.

11. Truong DT, Franzosa EA, Tickle TL, et al. MetaPhlAn2 for enhanced metagenomic taxonomic profiling. Nat Methods 2015; 12(10): 902–3.

12. Li D, Luo R, Liu CM, et al. MEGAHIT v1.0: A fast and scalable metagenome assembler driven by advanced methodologies and community practices. Methods 2016; 102: 3–11.

13. Hyatt D, Chen GL, Locascio PF, Land ML, Larimer FW, Hauser LJ. Prodigal: prokaryotic gene recognition and translation initiation site identification. BMC Bioinform 2010; 11: 119.

14. Li W, Godzik A. Cd-hit: a fast program for clustering and comparing large sets of protein or nucleotide sequences. Bioinformatics 2006; 22(13): 1658–9.

15. Pedersen HK, Forslund SK, Gudmundsdottir V, et al. A computational framework to integrate high-throughput ‘-omics’ datasets for the identification of potential mechanistic links. Nat Protoc 2018; 13(12): 2781–800.

16. Nielsen HB, Almeida M, Juncker AS, et al. Identification and assembly of genomes and genetic elements in complex metagenomic samples without using reference genomes. Nat Biotechnol 2014; 32(8): 822–8.

17. Aramaki T, Blanc-Mathieu R, Endo H, et al. KofamKOALA: KEGG Ortholog assignment based on profile HMM and adaptive score threshold. Bioinformatics 2020; 36(7): 2251–2.

18. de Oliveira GLV, Leite AZ, Higuchi BS, Gonzaga MI, Mariano VS. Intestinal dysbiosis and probiotic applications in autoimmune diseases. Immunology 2017; 152(1): 1–12.

19. Lugli GA, Tarracchini C, Alessandri G, et al. Decoding the genomic variability among members of the Bifidobacterium dentium species. Microorganisms 2020; 8(11): 1720.

20. Silvestre-Rangil J, Bagán L, Silvestre FJ, Bagán JV. Oral manifestations of rheumatoid arthritis. A cross-sectional study of 73 patients. Clin Oral Investig 2016; 20(9): 2575–80.

21. Marriott D, Stark D, Harkness J. Veillonella parvula discitis and secondary bacteremia: a rare infection complicating endoscopy and colonoscopy? J Clin Microbiol 2007; 45(2): 672–4.

22. Balakrishnan B, Luckey D, Taneja V. Autoimmunity-associated gut commensals modulate gut permeability and immunity in humanized mice. Mil Med 2019; 184(Suppl 1): 529–36.

23. Tito RY, Cypers H, Joossens M, et al. Brief report: Dialister as a microbial marker of disease activity in spondyloarthritis. Arthritis Rheumatol 2017; 69(1): 114–21.

24. Henrotin Y, Mathy M, Sanchez C, Lambert C. Chondroitin sulfate in the treatment of osteoarthritis: from in vitro studies to clinical recommendations. Ther Adv Musculoskelet Dis 2010; 2(6): 335–48.

25. Jura-Półtorak A, Komosinska-Vassev K, Kotulska A, et al. Alterations of plasma glycosaminoglycan profile in patients with rheumatoid arthritis in relation to disease activity. Clin Chim Acta 2014; 433: 20–7.

26. Shamdani S, Chantepie S, Flageollet C, et al. Heparan sulfate functions are altered in the osteoarthritic cartilage. Arthritis Res Ther 2020; 22(1): 283.

27. Severmann AC, Jochmann K, Feller K, et al. An altered heparan sulfate structure in the articular cartilage protects against osteoarthritis. Osteoarthr Carti 2020; 28(7): 977–87.

28. Carr AC, Maggini S. Vitamin C and immune function. Nutrients 2017; 9(11): 1211.

29. Abrams E, Sandson J. Effect of ascorbic acid on rheumatoid synovial fluid. Ann Rheum Dis 1964; 23(4): 295–9.

30. Abdelkarem HM, Fadda LH, El-Sayed EM, Radwan OK. Potential role of L-arginine and vitamin E against bone loss induced by nano-zinc oxide in rats. J Diet Suppl 2018; 15(3): 300–10.

31. Fiore CE, Pennisi P, Cutuli VM, Prato A, Messina R, Clementi G. L-arginine prevents bone loss and bone collagen breakdown in cyclosporin A-treated rats. Eur J Pharmacol 2000; 408(3): 323–6.

32. Manfredo Vieira S, Hiltensperger M, Kumar V, et al. Translocation of a gut pathobiont drives autoimmunity in mice and humans. Science 2018; 359(6380): 1156–61.

33. Scher JU, Sczesnak A, Longman RS, et al. Expansion of intestinal Prevotella copri correlates with enhanced susceptibility to arthritis. Elife 2013; 2: e01202.

34. Brown NA, Holt D, Webb M. The teratogenicity of methoxyacetic acid in the rat. Toxicol Lett 1984; 22(1): 93–100.

35. Miller RR, Carreon RE, Young JT, McKenna MJ. Toxicity of methoxyacetic acid in rats. Fundam Appl Toxicol 1982; 2(4): 158–60.

36. Sparks NR. The embryotoxic effects of harm reduction tobacco products on osteoblasts developing from human embryonic stem cells. University of California, Riverside 2018; chapter 4: 73–104.

37. Li P, Sundh D, Ji B, et al. Metabolic alterations in older women with low bone mineral density supplemented with Lactobacillus reuteri. JBMR Plus 2021; 5(4): e10478.

