## appendix for "Microbial dysbiosis and metabolic disorders promote rheumatoid arthritis across successive stages: a multi-omics cohort study"

### Supplementary Material

#### Supplementary Methods

##### The Cohort for Detection of Bacteria in the Joint Synovial Fluid

A total of 271 patients with RA of four distinct stages were recruited, including 52 RAS1, 66 RAS2, 67 RAS3, and 86 RAS4. RA patients were grouped into four RA stages according to the rheumatoid diagnostic score.[1] The patients met the aforementioned standard. Synovial- fluid samples were collected aseptically from knee joints during therapeutic aspiration. Synovial-fluid samples were deposited in sterile tubes on ice and homogenized within five minutes of collection. A tube filled with sterile phosphate-buffered saline (PBS) was left open throughout the procedure and subsequently processed in parallel with the samples as a negative control. The entire experiment was conducted in a completely sterile atmosphere. Each sample was immediately frozen and kept without heparin or hyaluronidase at –135 °C. For each patient, a total of 7ml synovial fluid was collected, of which 5ml was utilized for 16S rRNA gene sequencing, 1ml was used for bacteria isolation, and 1ml synovial fluid was prepared for scanning electron microscopy.

##### 16S rRNA Gene Sequencing and Processing

Bacterial DNA was extracted from 271 5ml synovial fluid samples. The tube containing PBS serves as environmental control. 86 synovial fluid samples from patients in RAS4 had bacteria DNA content (≥10ng) (Bacterial DNA Kit, TIANGEN) for bacteria 16S rRNA gene high-throughput sequencing. The V1/V2 hypervariable regions of the 16S ribosomal RNA gene were sequenced using the Illumina HiSeq platform. The 16S sequence paired-end data set was joined and quality filtered using the FLASH as previously described.[11] Sequence analysis was performed using the QIIME (version 1·9·1).[12] Chimeric sequences were removed using de novo chimera detection of USEARCH.[13] Sequences were clustered against the 2013 Greengenes (13_8 release) ribosomal database's 97% reference data set. Sequences that did not match any entries in this reference were subsequently clustered into de novo OTUs at 97% similarity with UCLUST.[13] Taxonomy was assigned to all OTUs using the RDP classifier[14] against Greengenes reference data set.

##### Isolation of Bacteria in the Joint Synovial Fluid

Due to that pH of joint synovial fluid was 7·6, we used Luria-Bertani (LB) broth medium with a pH of 7·6 for cultivation. The procedures were as follows: 1) sterilized LB medium (pH7·6) and inoculation equipment were placed in an anaerobic glove box (Ruskinn Concept 400) for deaeration one day in advance; 2) 1ml synovial fluid samples per stage of RA were used for bacterial cultivation. 3) The synovial fluid sample was serially diluted with sterilized water, plated onto LB (0·5% yeast extract, 1% tryptone, 1% sodium chloride) agar medium, and then incubated at 37°C for 72 h to obtain single colonies. Then, three colonies per plate were randomly selected and streaked three consecutive times on LB agar medium to obtain a pure culture, which was named isolate SF1 to SF9, respectively. To identify isolate SF1 to SF9, a partial fragment of 16S rDNA was amplified with the primer pair 27F (5'-AGAGTTTGATCCTGGCTCAG-3') and 1492R (5'-GGTTACCTTGTTACGACTT-3'), and then DNA sequencing was performed for preliminary identification.

**Scanning Electron Microscopy for Bacteria in the Joint Synovial Fluid**

The synovial fluid sample was filtered through the membrane (special membrane for flow cytometry) to remove solids and large particles (Filtrate A). Filtrate A was then filtered through a 0·45 μmum membrane to remove most human cells (Filtrate B). Filtrate B was then filtered through a 0·22 μmum filter membrane to enrich bacteria and subsequently washed with sterilized ultra-pure water to obtain Filtrate C. Filtrate C was soaked in 2·5% glutaraldehyde for 4 hours at room temperature. Filtrate C was then washed three times with 0·1 M PBS buffer and treated with 1% osmic acid for 4h. Filtrate C was then dehydrated in ethanol, vacuum dried by tert-butyl alcohol, coated with gold, and imaged with a scanning electron microscope (ZEISS Sigma 300).

**Gavage Experiments Using Ascorbate in the CIA Mouse Model**

Nine healthy seven-week-old DBA/1 mice weighing 20g were fed in an ultra-clean animal laboratory (SPF grade) with a humidity of 55% and a temperature of 26°C. CIA models were constructed and established as described before.[10] Nine mice were then divided into three groups (three mice per group), including normal DBA/1 mice and two groups of DBA/1 mice with CIA. Three-month gavage (0·3ml/d) to three groups of mice was conducted, including 1) 0·9% normal saline to normal DBA/1 mice, 2) 0·9% normal saline to DBA/1 mice with CIA, and 3) 100ng/ul ascorbate to DBA/1 mice with CIA. After three-month gavage, the mice plasma TNF-α level and the IL-6 level were tested using ELISA kit (mlbio, China). Mice were then killed and preserved in 4% formalin for two days. Micro-CT (QuantumGX, PerkinElmer, UnitedStates) was used to perform scanning and three-dimensional structural reconstruction of the joints. The settings were set to 209m, 90kV X-ray tube voltage, 160uA of the current, and 3 minutes of the scan time. The angle of the X-ray scan rotated 180 degrees. We have adhered to standards articulated in the Animal Research: Reporting of In Vivo Experiments (ARRIVE).

#### Supplementary Results

| Microbial species | RA (n=76) | | Healthy (n=27) | | Wilcox test | | Enrichment |
|---|---|---|---|---|---|---|---|
| | Mean | Std. Error | Mean | Std. Error | *p* | *q* | |
| *Bifidobacterium dentium* | 3.97E-04 | 1.42E-04 | 3.70E-05 | 3.70E-05 | 2.16E-03 | 3.43E-02 | RA |
| *Bacteroides uniformis* | 1.97E-02 | 5.05E-03 | 7.83E-02 | 2.30E-02 | 1.27E-04 | 1.24E-02 | Healthy |
| *Coprobacter fastidiosus* | 1.36E-04 | 4.72E-05 | 2.14E-03 | 1.41E-03 | 3.30E-03 | 3.94E-02 | Healthy |
| *Parabacteroides distasonis* | 3.50E-03 | 1.05E-03 | 4.97E-03 | 1.37E-03 | 2.45E-03 | 3.43E-02 | Healthy |
| *Streptococcus anginosus* | 3.00E-04 | 1.34E-04 | 0.00E+00 | 0.00E+00 | 4.17E-04 | 2.05E-02 | RA |
| *Streptococcus parasanguinis* | 6.46E-04 | 1.91E-04 | 3.92E-05 | 2.56E-05 | 1.08E-03 | 2.95E-02 | RA |
| *Clostridium hathewayi* | 8.54E-04 | 5.78E-04 | 9.63E-07 | 6.71E-07 | 1.58E-03 | 3.09E-02 | RA |
| *Veillonella atypica* | 3.27E-04 | 7.07E-05 | 6.54E-05 | 3.46E-05 | 4.02E-03 | 3.94E-02 | RA |
| *Escherichia coli* | 6.49E-02 | 1.43E-02 | 4.74E-03 | 1.97E-03 | 1.20E-03 | 2.95E-02 | RA |
| *Akkermansia muciniphila* | 2.53E-03 | 1.01E-03 | 3.43E-03 | 1.33E-03 | 3.72E-03 | 3.94E-02 | Healthy |

**Supplementary Table 1.** Significantly different microbial species between RA and healthy groups. Mean relative abundances of these species were showed. $p$ is produced using Mann-Whitney-Wilcoxon test, and $q$ is produced using Benjamini and Hochberg corrections. Results with $p < 0{\cdot}05$ and $q < 0{\cdot}05$ are displayed here. RA=rheumatoid arthritis.

| Microbial species | OA (n=19) | | Healthy (n=27) | | Wilcox test | | Enrichment |
|---|---|---|---|---|---|---|---|
| | Mean | Std. Error | Mean | Std. Error | *p* | *q* | |
| *Streptococcus anginosus* | 1.29E-04 | 5.94E-05 | 0.00E+00 | 0.00E+00 | 2.15E-03 | 3.52E-02 | OA |
| *Bacteroides cellulosilyticus* | 6.27E-05 | 6.01E-05 | 7.73E-03 | 4.77E-03 | 4.72E-04 | 1.54E-02 | Healthy |
| *Enterococcus faecium* | 3.80E-04 | 1.81E-04 | 0.00E+00 | 0.00E+00 | 7.94E-04 | 1.94E-02 | OA |
| *Ruminococcus torques* | 8.49E-03 | 2.79E-03 | 1.46E-03 | 3.78E-04 | 3.72E-04 | 1.54E-02 | OA |
| *Coprococcus catus* | 2.17E-03 | 7.19E-04 | 3.24E-04 | 1.43E-04 | 6.27E-05 | 6.14E-03 | OA |
| *Coprococcus comes* | 1.39E-03 | 3.27E-04 | 2.96E-04 | 8.87E-05 | 1.85E-03 | 3.52E-02 | OA |

**Supplementary Table 2.** Significantly different microbial species between OA and healthy groups. Mean relative abundances of these species were showed. $p$ is produced using Mann-Whitney-Wilcoxon test, and $q$ is produced using Benjamini and Hochberg corrections. Results with $p < 0{\cdot}05$ and $q < 0{\cdot}05$ are displayed here. OA=osteoarthritis.

| Microbial species | RAS1 (n=15) | | Healthy (n=27) | | Wilcox test | Enrichment |
|---|---|---|---|---|---|---|
| | Mean | Std. Error | Mean | Std. Error | | |
| *Streptococcus infantis* | 5.93E-06 | 5.63E-06 | 6.18E-05 | 2.76E-05 | 3.43E-03 | RAS1 |
| *Bacteroides ovatus* | 1.60E-02 | 8.91E-03 | 1.22E-03 | 4.45E-04 | 4.21E-03 | Healthy |
| *Atopobium parvulum* | 0.00E+00 | 0.00E+00 | 1.58E-05 | 8.60E-06 | 1.87E-02 | RAS1 |
| *Parasutterella excrementihominis* | 1.47E-03 | 6.70E-04 | 2.80E-04 | 2.62E-04 | 2.44E-02 | Healthy |
| *Granulicatella adiacens* | 6.59E-07 | 6.59E-07 | 3.08E-05 | 1.94E-05 | 2.51E-02 | RAS1 |
| *Turicibacter sanguinis* | 1.11E-06 | 1.11E-06 | 8.02E-05 | 4.57E-05 | 2.51E-02 | RAS1 |
| *Collinsella aerofaciens* | 3.56E-04 | 1.73E-04 | 2.00E-03 | 9.48E-04 | 4.34E-02 | RAS1 |

**Supplementary Table 3.** Significantly different microbial species between RAS1 and healthy groups. Mean relative abundances of these species were showed. $p$ is produced using Mann-Whitney-Wilcoxon test. Results with $p < 0{\cdot}05$ are displayed here. RAS1=the first stage of rheumatoid arthritis.

| Microbial species | RAS2 (n=21) | | Healthy (n=27) | | Wilcox test | Enrichment |
|---|---|---|---|---|---|---|
| | Mean | Std. Error | Mean | Std. Error | | |
| *Clostridium clostridioforme* | 1.12E-05 | 1.12E-05 | 3.94E-04 | 2.77E-04 | 1.95E-02 | RAS2 |
| *Bacteroides coprocola* | 7.47E-02 | 2.01E-02 | 8.26E-03 | 4.20E-03 | 3.57E-02 | Healthy |
| *Parabacteroides johnsonii* | 4.67E-04 | 2.35E-04 | 2.31E-06 | 2.31E-06 | 4.37E-02 | Healthy |
| *Scardovia inopinata* | 0.00E+00 | 0.00E+00 | 5.68E-06 | 3.95E-06 | 4.76E-02 | RAS2 |
| *Porphyromonas somerae* | 0.00E+00 | 0.00E+00 | 1.26E-05 | 9.64E-06 | 4.76E-02 | RAS2 |

**Supplementary Table 4.** Significantly different microbial species between RAS2 and healthy groups. Mean relative abundances of these species were showed. $p$ is produced using Mann-Whitney-Wilcoxon test. Results with $p < 0{\cdot}05$ are displayed here. RAS2=the second stage of rheumatoid arthritis.

| Microbial species | RAS3 (n=18) | | Healthy (n=27) | | Wilcox test | Enrichment |
|---|---|---|---|---|---|---|
| | Mean | Std. Error | Mean | Std. Error | | |
| *Parascardovia denticolens* | 0.00E+00 | 0.00E+00 | 1.25E-05 | 8.26E-06 | 1.21E-02 | RAS3 |
| *Lactobacillus gasseri* | 0.00E+00 | 0.00E+00 | 1.13E-04 | 6.15E-05 | 1.21E-02 | RAS3 |
| *Veillonella parvula* | 2.66E-04 | 8.53E-05 | 1.49E-03 | 6.11E-04 | 2.73E-02 | RAS3 |
| *Lactobacillus crispatus* | 0.00E+00 | 0.00E+00 | 4.82E-05 | 2.97E-05 | 3.21E-02 | RAS3 |
| *Lactobacillus oris* | 0.00E+00 | 0.00E+00 | 3.00E-05 | 1.84E-05 | 3.21E-02 | RAS3 |
| *Streptococcus mutans* | 0.00E+00 | 0.00E+00 | 3.72E-05 | 2.02E-05 | 3.21E-02 | RAS3 |
| *Dorea longicatena* | 9.42E-04 | 3.77E-04 | 3.21E-03 | 1.61E-03 | 3.74E-02 | RAS3 |
| *Enterobacter cloacae* | 4.13E-03 | 3.98E-03 | 4.51E-03 | 3.67E-03 | 3.75E-02 | RAS3 |

**Supplementary Table 5.** Significantly different microbial species between RAS3 and healthy groups. Mean relative abundances of these species were showed. $p$ is produced using Mann-Whitney-Wilcoxon test. Results with $p < 0{\cdot}05$ are displayed here. RAS3=the third stage of rheumatoid arthritis.

| Microbial species | RAS4 (n=22) | | Healthy (n=27) | | Wilcox test | Enrichment |
|---|---|---|---|---|---|---|
| | Mean | Std. Error | Mean | Std. Error | | |
| *Eggerthella lenta* | 1.56E-06 | 1.56E-06 | 1.11E-04 | 7.23E-05 | 1.82E-02 | RAS4 |
| *Bacteroides faecis* | 3.79E-03 | 1.47E-03 | 1.57E-04 | 1.02E-04 | 1.95E-02 | Healthy |
| *Bifidobacterium longum* | 1.01E-03 | 3.23E-04 | 1.05E-02 | 5.10E-03 | 2.25E-02 | RAS4 |
| *Lactococcus garvieae* | 0.00E+00 | 0.00E+00 | 7.46E-05 | 5.37E-05 | 2.35E-02 | RAS4 |
| *Bacteroides caccae* | 1.02E-02 | 3.41E-03 | 3.06E-03 | 1.03E-03 | 3.30E-02 | Healthy |

**Supplementary Table 6.** Significantly different microbial species between RAS4 and healthy groups. Mean relative abundances of these species were showed. $p$ is produced using Mann-Whitney-Wilcoxon test. Results with $p < 0{\cdot}05$ are displayed here. RAS4=the fourth stage of rheumatoid arthritis.

| Microbial species | OA (n=19) | | Healthy (n=27) | | Wilcox test | Enrichment |
|---|---|---|---|---|---|---|
| | Mean | Std. Error | Mean | Std. Error | | |
| *Coprococcus catus* | 3.24E-04 | 1.43E-04 | 2.17E-03 | 7.19E-04 | 6.27E-05 | OA |
| *Blautia obeum* | 1.26E-03 | 4.87E-04 | 2.91E-03 | 7.54E-04 | 4.77E-03 | OA |
| *Clostridium perfringens* | 0.00E+00 | 0.00E+00 | 1.18E-04 | 7.26E-05 | 3.71E-02 | OA |
| *Dialister invisus* | 8.17E-04 | 7.28E-04 | 2.22E-03 | 1.31E-03 | 4.10E-02 | OA |

**Supplementary Table 7.** Significantly different microbial species between OA and healthy groups. Mean relative abundances of these species were showed. $p$ is produced using Mann-Whitney-Wilcoxon test. Results with $p < 0{\cdot}05$ are displayed here. OA=osteoarthritis.

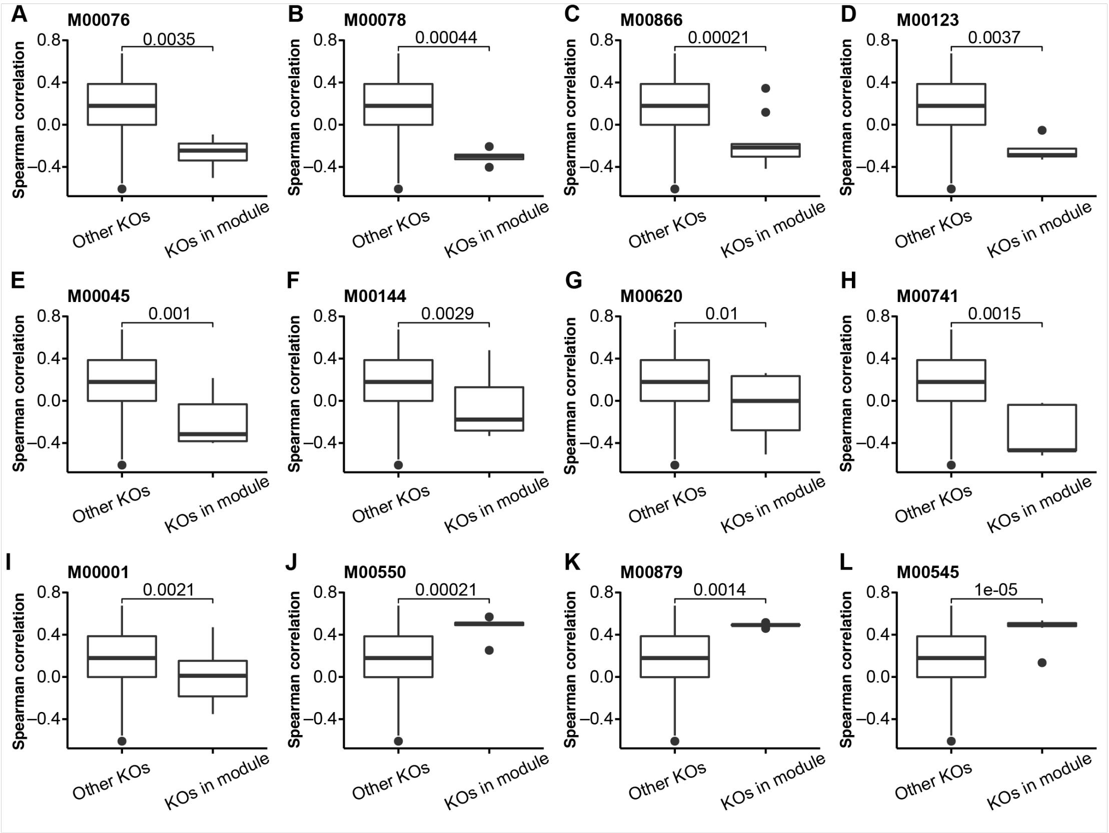


**Supplementary Figure 1.** KEGG modules that were identified as correlated with arthritis. The correlations of clinical phenotypes (Healthy=0, OA=1, RA=2) with KEGG modules were determined by if Spearman correlations of the phenotype with the abundances of KOs in the given KEGG module were significantly higher or lower (Mann–Whitney U-test FDR < 0·1) than with the abundances of all other KOs. Boxes represent the interquartile range between first and third quartiles and the line inside represents the median. Whiskers denote the lowest and highest values within 1·5 × interquartile range from the first and third quartiles, respectively. All the statistical significance was calculated by Mann-Whitney-Wilcoxon test.

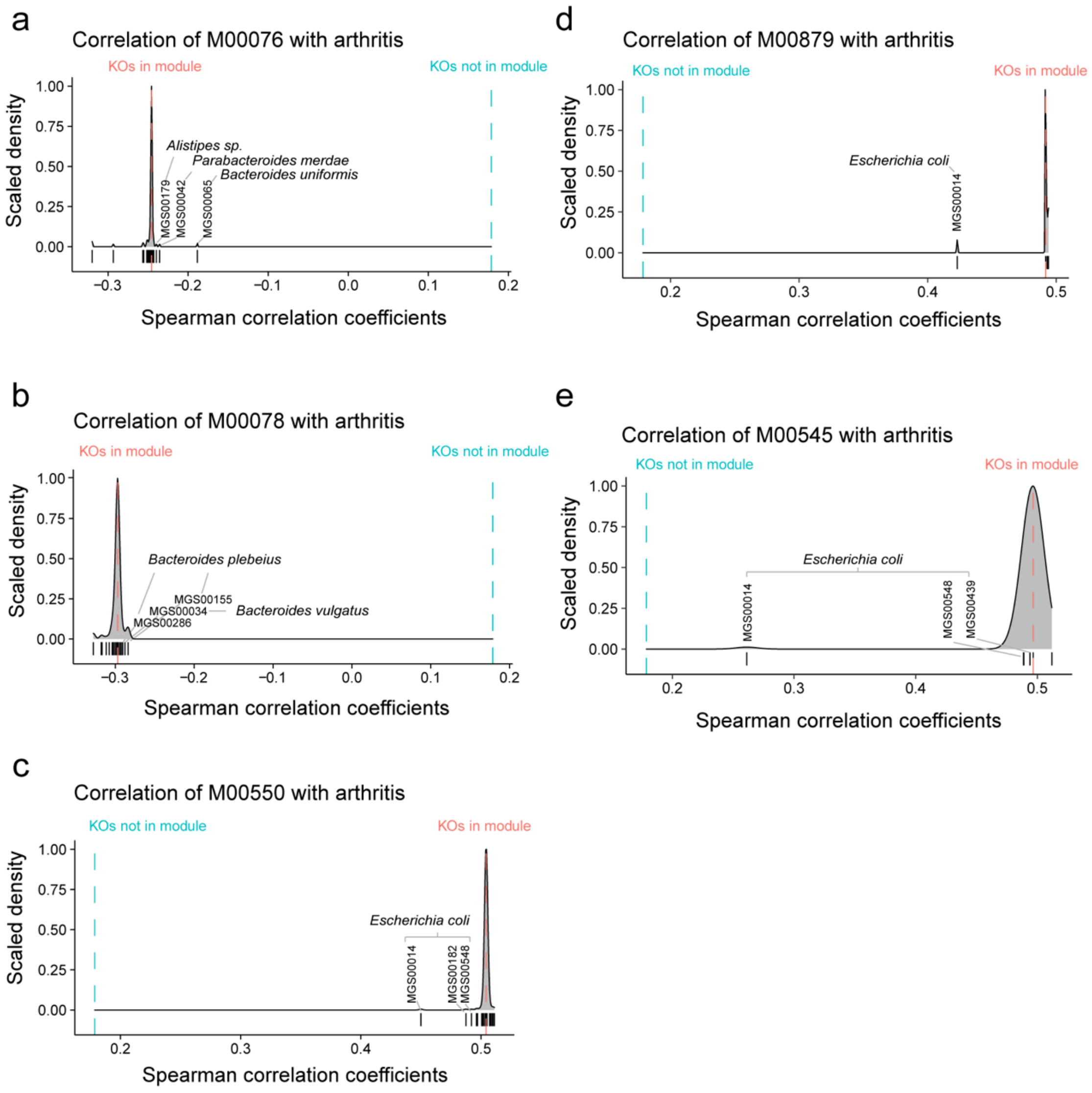


**Supplementary Figure 2.** The MGS that particularly contributed to the observed linkage between functional modules and arthritis. Dashed line represents the median SCC of the phenotypes (Healthy=0, OA=1, RA=2) with KOs in the specific module (red) and all other KOs (blue). Density plot shows the median SCC of the phenotypes with KOs in the specific module, when a given MGS (indicated by short vertical lines) has been excluded from the analysis. SCC=Spearman correlation coefficient. KO=KEGG ortholog. MGS=metagenomic species.

| Module | RF, IU/mL | IL-1β, pg/mL | IL-4, pg/mL | IL-8, pg/mL | IFN-γ, pg/mL | IL10, pg/mL | IL-12p70, pg/mL | IL13, pg/mL | IL-17, pg/mL | IL-2 , pg/mL | IL-6, pg/mL | TNF-α, pg/mL |
|---|---|---|---|---|---|---|---|---|---|---|---|---|
| **M00550** | | | | | | | | | | | | |
| *p* | 2.17E-01 | 5.44E-04 | 4.20E-03 | 7.45E-04 | 3.13E-03 | 1.55E-03 | 3.47E-03 | 8.58E-02 | 1.04E-01 | 9.37E-04 | 1.12E-03 | 6.59E-04 |
| *q* | 5.47E-01 | 3.70E-02 | 9.67E-02 | 7.07E-02 | 1.06E-01 | 1.05E-01 | 9.31E-02 | 4.21E-01 | 4.21E-01 | 6.37E-02 | 6.11E-02 | 5.04E-02 |
| M00879 | | | | | | | | | | | | |
| *p* | 1.35E-03 | 1.26E-02 | 2.43E-01 | 2.01E-02 | 7.60E-02 | 1.62E-02 | 1.72E-01 | 1.90E-02 | 8.29E-01 | 1.46E-02 | 1.25E-01 | 3.18E-02 |
| *q* | 4.72E-02 | 1.96E-01 | 5.50E-01 | 2.97E-01 | 5.19E-01 | 2.46E-01 | 4.58E-01 | 2.15E-01 | 9.15E-01 | 2.33E-01 | 4.67E-01 | 3.60E-01 |
| M00545 | | | | | | | | | | | | |
| *p* | 1.30E-04 | 2.87E-05 | 1.77E-03 | 2.46E-04 | 3.92E-04 | 4.10E-05 | 2.61E-03 | 1.63E-05 | 7.77E-01 | 7.40E-05 | 6.12E-03 | 6.97E-05 |
| *q* | 1.18E-02 | 3.91E-03 | 6.07E-02 | 6.69E-02 | 3.56E-02 | 5.57E-03 | 8.88E-02 | 4.41E-03 | 8.95E-01 | 1.01E-02 | 1.51E-01 | 9.47E-03 |

**Supplementary Table 8.** Correlations of KEGG modules with rheumatoid factor and plasma cytokines. The correlations of cytokine level with KEGG modules were determined by if Spearman correlations of the cytokine level with the abundances of KOs in the given KEGG module were significantly higher or lower than with the abundances of all other KOs. *p* is produced using Mann-Whitney-Wilcoxon test, and *q* is produced using Benjamini and Hochberg corrections. RF=rheumatoid factor. IL=interleukin. TNF=tumour necrosis factor. IFN=Interferon. M00550=ascorbate degradation. M00879= arginine succinyltransferase pathway. M00545=trans-cinnamate degradation.

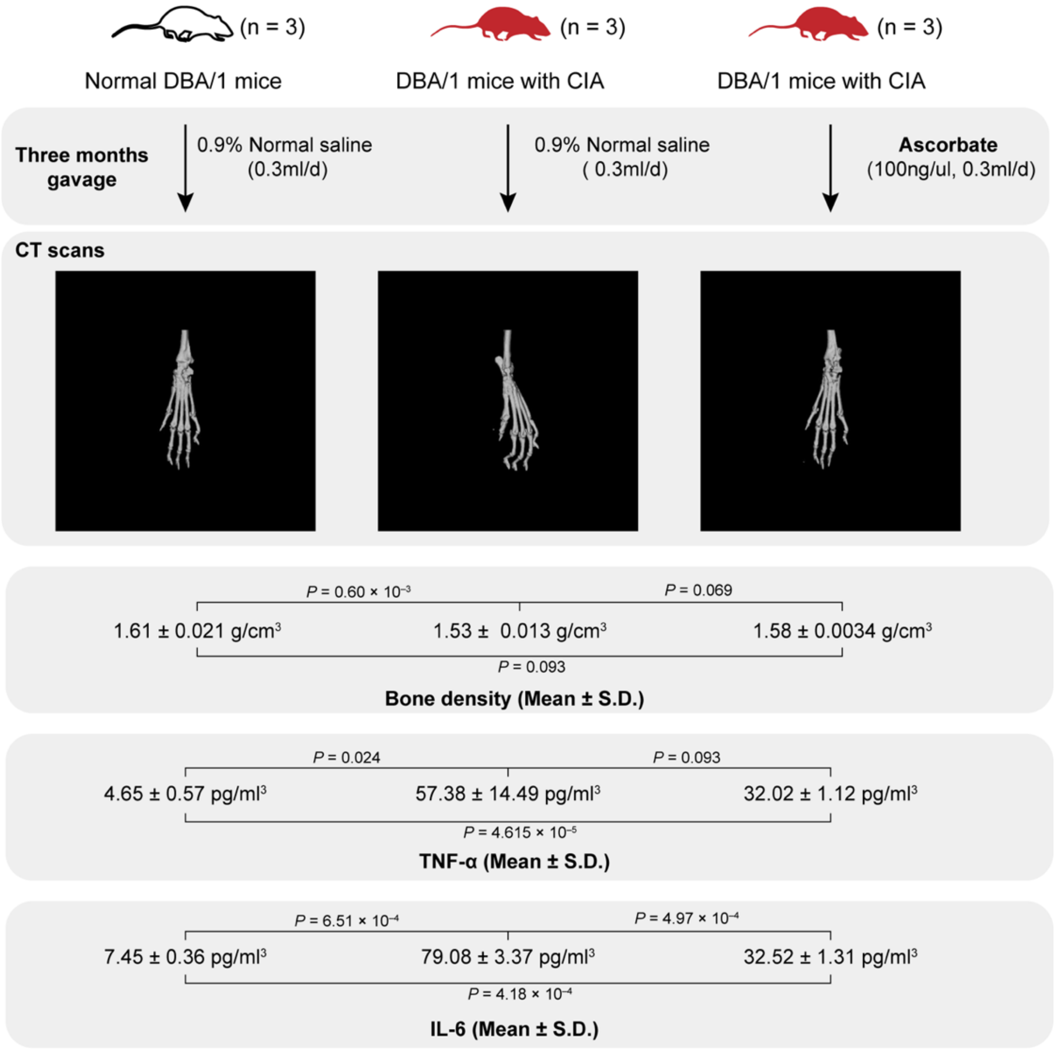


**Supplementary Figure 3** Ascorbate ameliorate collagen-induced arthritis in mice model. Three groups of mice (three mice per group) were used, including normal DBA/1 mice and two groups of DBA/1 mice with CIA. After three-month gavage, the CT scans, bone density, plasma TNF-α level, and plasma IL-6 level were examined. The statistical test is performed using t-test.

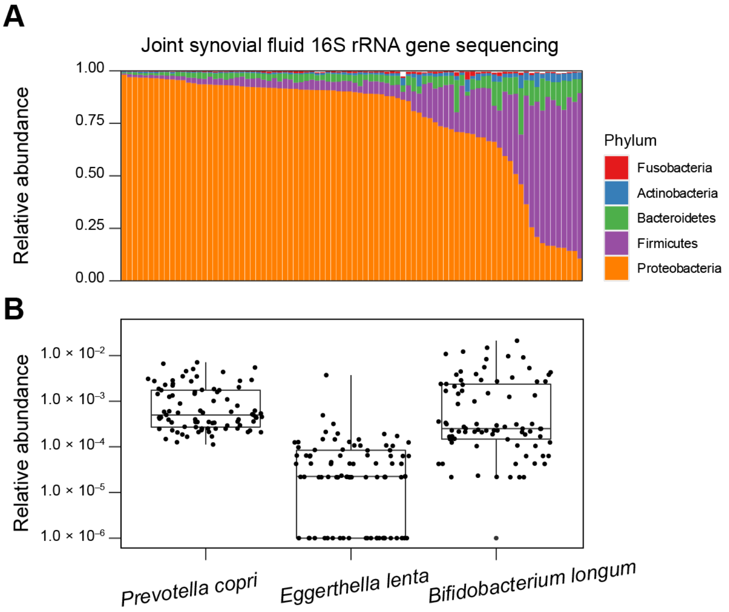


**Supplementary Figure 4** Microbial composition in the joint synovial fluid of patients in RAS4. (**A**) Relative abundances of the five most abundant phyla in the joint synovial fluid of patients in RAS4. (**B**) Relative abundances of the three microorganisms in the joint synovial fluid of patients in RAS4. The boxes represent 25th–75th percentiles, black lines indicate the median and whiskers extend to the maximum and minimum values within 1.5 × the interquartile range. RAS4=the fourth stage of rheumatoid arthritis.

| Sample | Sequences | Taxonomy |
| --- | --- | --- |
| SF1, SF3, SF4, SF8, SF9 | GCTGGCTCCTTACGGTTACCTCACGGACTTCGGGTGTTACCAGCTCTCATGGTGTGACGGGC<br>GGTGTGTACAAGGCCCGGGAACGTATTCACCGCGACATTCTGATTCGCGATTACTAGCAAC<br>TCCAGCTTCATGTAGGCGAGTTGCAGCCTACAATCCGAACTGAGATAGGTTTTATAAGTTT<br>TGCTCCACCTCACGGTCTTGCGTCTTATTGTACCTACCATTGTAGCACGTGTGTAGCCCTGGA<br>CATAAGGGGCATGATGATTTGACGTCATCCCCACCTTCCTCCTGGTTACCCAGGCAGTCTCA<br>TTAGAGTGCTCAACTTAATGGTAGCAACTAATAACAAGGGTTGCGCTCGTTGCAGGACTTA<br>ACCTAACATCTCACGACACGAGCTGACGACAACCATGCACCACCTGTCTCCTTGCCCCGAAG<br>GGCTTCACCTATCTCTAGGCTATGCAAGGGATGTCAAGTCCAGGTAAGGTTCTTCGCGTTGC<br>TTCGAATTAAACCACATGCTCCGCTGCTTGTGCGGGCCCCCGTCAATTCCTTTGAGTTTAAT<br>CTTGCGACCGTACTCCCCAGGCGGGATACTTATTGTGTTAACTGCGGCACAGGGGGAGTTG<br>ATACCCCCTACACCTAGTATCCATCGTTTACGGCGTGGACTACCAGGGTATCTAATCCTGTT<br>TGCTACCCACGCTTTCGTGCCTCAGCGTCAGTTACAGTCCAGAAAGCCGCCTTCGCCACTGG<br>TGTTCTTCCTAATCTCTACGCATTTCACCGCTACACTAGGAATTCCGCTTTCCTCTCCTGCACT<br>CTAGATATCCAGTTTGGAATGCAGCCCCCAGGTTAAGCCCGGGGATTTCACATCCCACTTAA<br>ACATCCGCCTACGCACCCTTTACGCCCAGTAAATCCGGACAACGCTCGCCACCTACGTATTA<br>CCGCGGCTGCTGGCACGTAGTTAGCCGTGGCTTCCTCCTCTGGTACCGTCATTATCGTCCCAG<br>AAAACAGGGCTTTACAATCCGAAGACCTTCATCACCCACGCGGCGTTGCTGCGTCAGGGTT<br>TCCCCCATTGCGCAATATTCCCCACTGCTGCCTCCCGTAGGAGTCTGGACCGTGTCTCAGTTC<br>CAATGTGGCCGATCACCCTCTCAGGTCGGCTACGCATCGTTGCCTTGGTAAGCCGTTACCTT<br>ACCAACTAGCTAATGCGCCGCGGGTCCATCTCAAAGCAATAAATCTTTGATAAGAAAATCA<br>TGCGATTCTCTTATGTTATGCGGTATTAATCTTCCTTTCGGAAGGCTATCCCCCACTTTGAGG<br>CAGGTTACCCACGTGTTACTCACCCGTCCGCCGCTAATCCACTTCCCGAAGGAAGCTTCATC<br>GCTCGACTTGCATGTGTTAAGCACGCCGCCAGCGTTCGTCCTGAGCC | 1.Clostridium sporogenes gene for 16S ribosomal RNA, partial sequence, strain JCM 7841<br>2.Clostridium sporogenes strain DSM 29422 |
| SF2, SF6, SF7 | GGGGCCTGCGCAGCCTATACATGCAAGTCGAACGCTTTTTCTTTCACCGGAGCTTGCTCCAC<br>CGAAAGAAAAAGAGTGGCGAACGGGTGAGTAACACGTGGGTAACCTGCCCATCAGAAGG<br>GGATAACACTTGGAAACAGGTGCTAATACCGTATAACACTATTTTCCGCATGGAAGAAAG<br>TTGAAAGGCGCTTTTGCGTCACTGATGGATGGACCCGCGGTGCATTAGCTAGTTGGTGAGG<br>TAACGGCTCACCAAGGCCACGATGCATAGCCGACCTGAGAGGGTGATCGGCCACACTGGG<br>ACTGAGACACGGCCCAGACTCCTACGGGAGGCAGCAGTAGGGAATCTTCGGCAATGGACG<br>AAAGTCTGACCGAGCAACGCCGCGTGAGTGAAGAAGGTTTTCGGATCGTAAAACTCTGTT<br>GTTAGAGAAGAACAAGGATGAGAGTAGAACGTTCATCCCTTGACGGTATCTAACCAGAAA<br>GCCACGGCTAACTACGTGCCAGCAGCCGCGGTAATACGTAGGTGGCAAGCGTTGTCCGGAT<br>TTATTGGGCGTAAAGCGAGCGCAGGCGGTTTCTTAAGTCTGATGTGAAAGCCCCCGGCTCA<br>ACCGGGGAGGGTCATTGGAAACTGGGAGACTTGAGTGCAGAAGAGGAGAGTGGAATTCC<br>ATGTGTAGCGGTGAAATGCGTAGATATATGGAGGAACACCAGTGGCGAAGGCGGCTCTCT<br>GGTCTGTAACTGACGCTGAGGCTCGAAAGCGTGGGGAGCGAACAGGATTAGATACCCTGG<br>TAGTCCACGCCGTAAACGATGAGTGCTAAGTGTTGGAGGGTTTCCGCCCTTCAGTGCTGCA<br>GCAAACGCATTAAGCACTCCGCCTGGGGAGTACGACCGCAAGGTTGAAACTCAAAGGAAT<br>TGACGGGGGCCCGCACAAGCGGTGGAGCATGTGGTTTAATTCGAAGCAACGCGAAGAACC<br>TTACCAGGTCTTGACATCCTTTGACCACTCTAGAGATAGAGCTTCCCCTTCGGGGGCAAAGT<br>GACAGGTGGTGCATGGTTGTCGTCAGCTCGTGTCGTGAGATGTTGGGTTAAGTCCCGCAAC<br>GAGCGCAACCCTTATTGTTAGTTGCCATCATTTAGTTGGGCACTCTAGCGAGACTGCCGGTG<br>ACAAACCGGAGGAAGGTGGGGATGACGTCAAATCATCATGCCCCTTATGACCTGGGCTAC<br>ACACGTGCTACAATGGGAAGTACAACGAGTTGCGAAGTCGCGAGGCTAAGCTAATCTCTT<br>AAAGCTTCTCTCAGTTCGGATTGTAGGCTGCAACTCGCCTACATGAAGCCGGAATCGCTAG<br>TAATCGCGGATCAGCACGCCGCGGTGAATACGTTCCCGGGCCTTGTACACACCGCCCGTCAC<br>ACCACGAGAGTTTGTAACACCCGAAGTCGGTGAGGTAACCTTTTGGAGCCAGCCGCCTAAG<br>TGGAAAAGGAAGT | Enterococcus sp. w17 16S ribosomal RNA gene, partial sequence |
| SF5 | GGGGCTGCGCGGCTACACTGCAGTCGAACGCTTCTTTCCTCCAGCTTGCTTGCTGTCTGACG<br>AGTGGCGGATGGTGAGTAGTGTATGGGAAACTGCCCGATGGAAGGGGATACCTACTGGAC<br>ACGGTAGCTAATACCGCATAACGTCTTGGACCATTATGGGGGACCTTCGGGCCTCTTGCCAT<br>CGGATGCGTGTCATGGGATTATGGACCCGGTGGGGTAACGGCTCACCTAGGCGACGATCCC<br>TAGATGGTCTGAATGATGACCAGCCATGCTGGGGCTATCACACCCACCAGACTCCAGCGGG<br>AGGCAGCACTGGGGAATATGGCACGATGGGGGCAAGCCTGATGCAGCACTGCCGGGTGTA<br>TGAAGAAGGCCTTCTGGTTGAAAAGTGTTTTCAGCGAGGATGAAGGTGTTGTGGTTAATA<br>ACCAAGGAAATTGACGTTACTCGCTCCCCTGACACCGGCTAACTCCGTGCAACGGCCGCTAC<br>ATACGGAGGGTGCAAGCGTTAATCGGAATTACTGCGCGTAAAGGACAATTGGGGGTCTGT<br>CAAGTCGGATGTGAAATTCCCGGGGATACCTGGGAACTGCATCCGAAACTGGAGGGCTTT<br>GTCTTGTGGAGGACGGTAGAATTCCAGGTGTAACGGTAATTTGCGGAGAGATCTGGAGGA<br>ATACCGATGGCGAAGGAGCCCCCCTGGACAAAGACTGACCCCTCAGGTGCTAACTCATGCT<br>GAGCCCCAGGATTAGATACCCTGATAGTACACGCCGTACACGATGTCCACTTGCAGGTTGA<br>GCACTTGAAGCGTGGCTACGGGAGCTAACGCGTTAGTCGACGACTGGAGAGTACCGCCGC<br>GAGGGTAAAACTCAATGAATGACGGGGTCCATACAAGCGGCGGAGCATGTGTTACATTTC<br>GATGCATCGCGTAGAATCTACCTACTCCTGACATCCCCATCAGGTTCTCAGAATTCCTTGGC<br>GGCCTGCGGGAACTCCTATACAGTGCGGGACAGATTGACGAGGGTCCATGTGTGACTTAA<br>GAGGCAGTCAGCCAACCCTGGAACTGAGACCTGTCAGACTCCTACGAGCAGCAGTGCGATC<br>ATCGCCATGACGACAGTCTGACTGCAGCCATGACGCGTGATTGCAGAGGCATCAGAGTTGT<br>AAAGTACGTCATCGAGAGACGTGATCGTCGATTATTAGCAGCGTGCCACTGACGTACTCAG<br>CAGAAGAAGCACCGGCTAACTACCGTGCCAGCAGCCGCGGTAATACGGAGGGGTGCAAGC<br>GTGTAATCGGAATTATTGGGCGTAAAGCGACACGCAGGCGGTCTGTCAAGTCGGATGTGA<br>AATCCCCCGGGCTCAACCGTGGGAGGGTGCATCTGAAACTGGCAGACTTAAAGTCTGATGG<br>AGGAGAGATGGGAATTCCATGTGTAGCGGGTGAAAATGCGTAGATAATATGGAAGGAAC<br>ACCAGTGGGCGAAGGGCGGGTCTCTTGGTACATAGACTGACGCTTCGGCTCGGAAAGCGT<br>GGGGGAAGCAAACACGGATTAGATACCCCTGTTATTCCCGCCCGTAACGGATGTCGACATG<br>GAGTGTGGTGGGCCTTCACCTGTGGCTTCCGGAGCTAACGCATTAAGTCACTCCGCCTGGG<br>GGAGTACGGCCGCAAGGTTAAAACTCAAAGGAATTGACGGGGGGCCCGCACAAGCGGTG<br>GAGCATGTGGTTTAATTCGATGCAACGCGAAGAACCTTACCACTCTTGACATCCAGAGACC<br>TAGCAGAGATAGAGCTTTCCTTCGGGAACTCAAGGACAGGGTGGGCATGGCGGTTTCTCAT<br>CGTGTTGTGAAATGTTGTATTATGTCTCGCACCGAGAGCACCCCTTATCCTTAGTGCCATCA<br>TTCGGTCGGCAACTCAAAGGAGACTCCCAGGGCTAAACTGGAGGAAGGTGGGGATGACGT<br>CAAGTCATCATGGCCCTTACGAGTAGGGCTCCACACGTGTTCCAATGGAAGATACAAAGAG<br>AAGCAACCCCGGGAGTCCAAGCAATCTTCATAAGCTTCTTTCAATTCCGGATGCAGGTTGC<br>AAATTGCCTTCATGAAGCTGGAAATCGTTAGTAATGGGGATCGCACGCCCCCGGTGAATAG<br>TTTCCCGGCCTTTACCCCCCCCCCGTCACCACCGAGAATTGGAACCCAAGAAGTTGGGGAGG<br>TAACCGTAAGGAGCCAGCTCCTAAGTGAACGTTGT | Citrobacter sp. 86.6 16S ribosomal RNA gene, partial sequence |

**Supplementary Table 8.** 16S rRNA gene sequencing of the isolated single colony from the joint synovial fluid of patients in the fourth stage of rheumatoid arthritis.

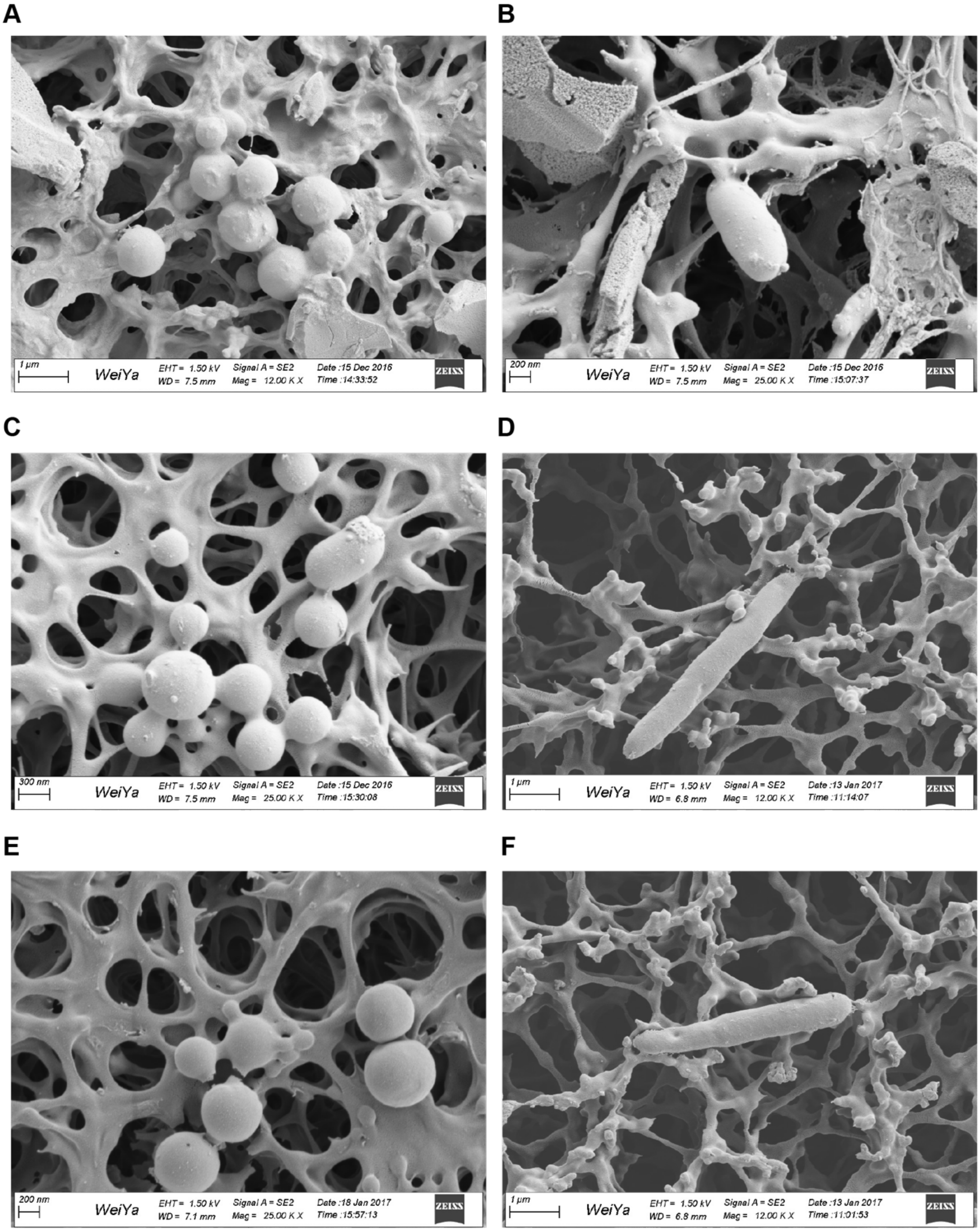


**Supplementary Figure 5** Scanning electron microscopy of the joint synovial fluid.
